# Aging, HIV Status, and Sex Differentially Influence Antiretroviral Therapy Disposition Pathways in Rectosigmoid Tissue and Peripheral Blood Mononuclear Cells

**DOI:** 10.64898/2026.11.05.764744

**Authors:** Hannah N. Wilkins, Cory J. White, Ruohui Zheng, Labib A. Mamun, Colten D Eberhard, Edward J. Fuchs, Jennifer Hoffmann, Ekta Gupta, Benjamin C. Orsburn, Craig W. Hendrix, Namandje N. Bumpus, Dionna W. Williams

## Abstract

People living with human immunodeficiency virus (HIV) are increasingly surviving into older age due to the effectiveness of antiretroviral (ARV) therapy (ART), creating a growing population in whom age-related changes may influence treatment outcomes. Although aging is known to affect drug disposition, the molecular mechanisms by which aging alters ARV pharmacology remain incompletely understood, particularly within tissue compartments, including the rectosigmoid region of the colon, that serve as sites of HIV transmission and persistence. To address this knowledge gap, we used proteomics by liquid chromatography-mass spectrometry (LC-MS) to evaluate the impact of age and HIV status on proteins and pathways relevant to ARV disposition in peripheral blood mononuclear cells (PBMC) and rectosigmoid colon tissue obtained from young (18-50) HIV-seronegative adults, older (65-80) HIV-seronegative adults, and older virally suppressed people living with HIV. We found that aging and HIV infection differentially influenced proteins predominantly within nucleos(t)ide reverse transcriptase inhibitor (NRTI) pathways, in rectosigmoid tissue and PBMC. HIV- associated effects to the proteome were more prominent in PBMC, whereas aging produced broader alterations in ARV disposition-related proteins in rectosigmoid tissue, including proteins involved in drug transport, intracellular drug activation, dephosphorylation, and nucleotide homeostasis. Several age-associated changes were further influenced by biologic sex, suggesting that aging may not affect ARV disposition uniformly across populations. To place these findings within a broader context, pathway analyses were performed that identified extensive tissue-specific remodeling associated with aging and HIV infection involving drug disposition, immune regulation, cellular metabolism, and disease-related pathways. Notably, aging was associated with alterations in pathways relevant to nucleotide metabolism and gastrointestinal tissue function, suggesting that processes beyond traditional drug transport and metabolism pathways may contribute to variability in ARV disposition with advancing age. Together, these findings identify aging as an important modifier of pathways that regulate ARV disposition within compartments relevant to HIV transmission and persistence. While the clinical consequences of our findings are unknown, these data provide a mechanistic foundation for future studies investigating whether age-associated changes influence the effectiveness of ART and HIV pre-exposure prophylaxis in older adults.

**Significance Statement:** Adults over 50 years of age comprise a rapidly growing proportion of both HIV treatment and prevention populations, yet the biological effects of aging on antiretroviral drug disposition remain poorly understood. We demonstrate that aging, sex, and HIV status differentially alter proteins and pathways involved in antiretroviral transport, activation, and nucleotide homeostasis, with the most pronounced effects occurring in colorectal tissue, providing a mechanistic foundation for future studies of HIV therapeutics in aging populations.

**Visual Abstract:** 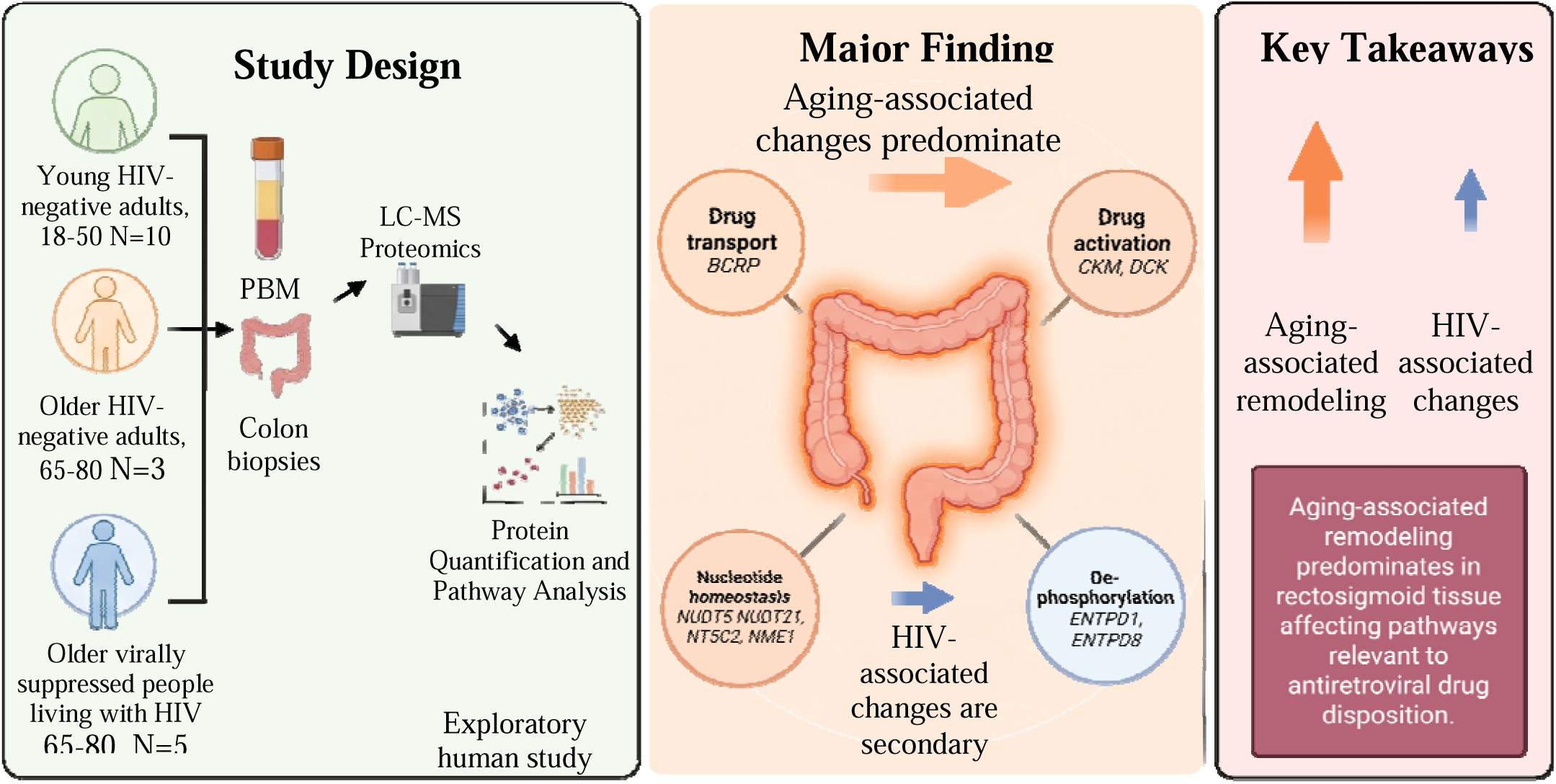

## Introduction

The prevalence of human immunodeficiency virus (HIV) has increased substantially in older adults (>50 years of age), which is largely due to the success of antiretroviral (ARV) therapies (ART) in extending life expectancy and decreasing mortality in people living with HIV (PLWH) ^1^. Indeed, approximately half of all PLWH in the United States are over the age of 50 ^1,2^. To this end, with proper adherence and access to ART, HIV has transitioned from an acute, life-limiting condition to a chronic, manageable infection. Aging of the HIV population has also occurred because of a shift in the demographics of new infections with ∼16-20% of new HIV diagnoses occur among individuals >50, highlighting that older adults comprise an important at-risk population ^3,4^.

Clinical care of aging individuals requires unique pharmacologic considerations to optimize ART efficacy. First, high medication burden to manage comorbid conditions is common in older PLWH. The concurrent use of multiple medications, termed polypharmacy, increases the potential for adverse drug-drug interactions and is associated with several adverse outcomes, including slow gait speed, recurrent falls, increased rates of hospitalization, and cognitive decline ^5–8^. Changes in immune responses during aging must also be considered because of age-related delays in immune reconstitution following ART initiation and immune system decline, as well as the cumulative effects of living with HIV-induced chronic inflammation and the premature aging that occurs during HIV, termed “inflammaging”, that leads to heightened inflammatory states ^9,10^. Finally, aging PLWH with cognitive impairment may have suboptimal ART adherence that poses major risks for viral suppression ^11,12^.

To this end, it is increasingly important to understand how aging influences ARV pharmacokinetics and pharmacodynamics. Aging alters renal clearance, hepatic blood flow, and liver mass, processes that may affect systemic ARV disposition ^13^. Consistent with this premise, older PLWH exhibit altered plasma and intracellular concentrations of tenofovir (TFV) and emtricitabine (FTC), two nucleotide reverse transcriptase inhibitors (NRTIs) that form the backbone of most ART regimens that are also widely used for HIV pre- exposure prophylaxis (PrEP) ^14,15^. These differences have been attributed to age-related changes that modify drug elimination, pharmacologic response, and the time required to achieve steady-state concentrations ^16,17^. However, current understanding remains incomplete because most studies have focused on plasma and peripheral blood mononuclear cells (PBMC), which may not reflect drug disposition within tissue reservoirs where HIV persists, including the colorectal tissue of the gastrointestinal (GI) tract ^18,19^.

This limitation is particularly important because intracellular ARV concentrations are determined by a complex network of proteins, including drug transporters, prodrug releasing enzymes, phase I and II metabolizing enzymes, activating kinases, dephosphorylating enzymes, and nucleotide homeostasis regulators that control endogenous nucleotide pools and NRTI disposition. These proteins collectively govern the activation, inactivation, and persistence of intracellular ARV metabolites and exhibit distinct tissue-specific expression patterns. Importantly, many are also susceptible to aging-related alterations ^15^. For example, hepatic expression of the transporter breast cancer resistance protein (BCRP) decreases with aging, whereas intestinal BCRP ^20^, P-glycoprotein (P-gp), multidrug resistance protein (MRP)-1 (MRP1), and MRP4 ^21^ expression may increase. Similarly, activity of creatine kinases (CK), isoenzymes involved in TFV activation, declines in aging brain and skeletal muscle ^22,23^. Together, these findings suggest that aging may differentially alter ARV disposition across tissues, yet the impact of aging on NRTI metabolic pathways within colorectal tissue remains largely unexplored. Because the GI tract represents both a major site of HIV transmission and a persistent viral reservoir, this knowledge gap limits our ability to understand how aging influences local ARV activity. Defining age-associated changes in proteins that regulate intracellular ARV concentrations may therefore provide important insight into ART efficacy in older adults that are HIV positive or at risk of HIV transmission and identify biologic factors that contribute to variability in treatment outcomes.

Here, we address this gap by evaluating the impact of age and HIV status on the relative abundance of ARV-relevant proteins in PBMC and rectosigmoid colon tissue, two compartments with distinct biologic roles in HIV infection. Using proteomics by liquid chromatography-mass spectrometry (LC-MS), we measured the relative abundance of drug transporters, phase I/II metabolizing enzymes, activating kinases, phosphatases, nucleotidases, nucleoside hydrolases, and other nucleotide homeostasis proteins in young HIV-seronegative adults (18-50 years, n=10), older HIV-seronegative adults (65-80 years, n=3), and older virally suppressed PLWH (65-80 years, n=5) (**Table 1**). We found that aging and HIV status altered numerous ARV-disposition related proteins, with the most pronounced effects occurring in rectosigmoid tissue. While HIV-associated changes predominated in PBMC, aging substantially affected colonic proteins involved in drug transport (BCRP), prodrug activation (CKM, DCK), dephosphorylation (ENTPD1, ENTPD8), and nucleotide homeostasis (NUDT5, NUDT21, NT5C2, NUDT16L1, and NUDT1). We further identified sex-dependent effects on several ARV-disposition related proteins and observed broad age- and HIV-associated alterations in pathways related to drug metabolism, immunity, energy metabolism, and disease pathology. Together, these findings demonstrate that age, sex, and HIV status differentially influence molecular determinants of ARV disposition in circulating PBMC and rectosigmoidal colonic tissue, highlighting the potential need for age-informed strategies to optimize ART efficacy. Given the challenges associated with recruiting older participants for paired blood and colorectal tissue sampling, this study was conducted as an exploratory, hypothesis-generating investigation and enrolled all eligible participants available during the study period. Although not powered to assess clinical outcomes, the novel age-, sex-, and HIV-associated differences identified in key ARV disposition pathways provide a mechanistic foundation for future studies designed to determine whether these proteomic changes influence protein function and treatment efficacy in aging populations across HIV status and sex. As PLWH age and the median age of HIV transmission increases, our findings identify molecular pathways that may influence ARV disposition in older adults and provide a rationale for future clinical studies, with larger sample sizes, evaluating whether treatment strategies developed in younger populations remain optimally effective across the aging spectrum.

**Table 1:** Participant demographics.

| Demographic (Age Range) | **N** | **Male, N (%)** | **Female, N (%)** | **African American, N (%)** | **Asian, N (%)** | **White, N (%)** |
|---|---|---|---|---|---|---|
| HIV-uninfected, Young (18-50) | 10 | 6 (60) | 4 (40) | 2 (15) | 1 (7) | 10 (76) |
| HIV-uninfected, Old (65-80) | 3 | 1 (33) | 2 (67) |  |  |  |
| HIV-positive, Old (65-80) | 5 | 5 (100) | 0 (0) | 3 (60) | 0 (0) | 2 (40) |

## Materials and Methods

### Clinical Study Visits

All prospective research participants provided written informed consent prior to study screening for eligibility. The study was approved for human subjects research by the Johns Hopkins Medicine Institutional Review Board (IRB00307426) and registered on clinicaltrials.gov (NCT06132386).

For prospective research participants not scheduled for routine colon cancer colonoscopic screening, a separate screening visit began with review and signing of the approved informed consent form followed by physical exam where height, weight, blood pressure and heart rate were measured. Rectal swabs were collected for gonorrhea and chlamydia testing. Blood was collected to evaluate plasma chemistry (including liver and kidney function), hematology, coagulation, and syphilis and HIV infection. For HIV negative participants (Control Cohort A), screening confirmed no known risk for HIV exposure or a documented negative HIV-1/HIV-2 Ag/Ab test in the past 3 months (HIV risk, but not on PrEP). For participants infected with HIV (ART Cohort B), screening included review of prior test result documentation confirming virologically suppressed HIV for at least 6 months prior to screening as well as a history of taking a tenofovir disoproxil fumarate (TDF) or tenofovir alafenamide (TAF) containing regimen to treat HIV. Participants were excluded for inflammatory bowel disease or other active inflammatory conditions of the gastrointestinal tract, as well as investigational, immunomodulatory or rectally-administered medications. Following screening, eligible research participants completed a single study sampling visit which included interim medical history, vital signs, directed physical exam, confirmation of overnight fast and self administration of a cleansing rectal enema prior to arriving in the research and endoscopy clinic, blood collection for proteomic and pharmacokinetic analysis, and a flexible sigmoidoscopy or colonoscopy for rectosigmoid collection. Approximately 50 mL of blood was acquired for proteomic testing and future pharmacokinetic testing. 10 rectosigmoid biopsies were collected from the rectum within 10-20 cm of the anal verge by flexible sigmoidoscopy or colonoscopy using single-use 240 cm radial jaw biopsy forceps (Boston Scientific, Marlborough, MA) threaded through a flexible sigmoidoscope or colonoscope.

For participants already scheduled for a colonoscopy for routine preventive health colon cancer screening, the screening visit consisted of only the informed consent process, history, blood HIV and rectal GC/CT testing, and could occur on the same day and location as the sampling visit, immediately prior to the scheduled colonoscopy. Participant visits took place at the Johns Hopkins Hospital Drug Development Unit and Endoscopy Suite (Baltimore, MD) or Johns Hopkins Gastroenterology and Hepatology (Columbia, MD).

### Rectosigmoid Biopsy Processing

Following collection, rectosigmoid colon biopsies were placed on ice and transported to the laboratory where they were weighed, flash frozen, and stored at -80°C until LC-MS analysis. Whole blood was immediately placed on ice and processed within 2 hours following collection for PBMC isolation. TFV and TFV diphosphate (TFV-DP) were quantified in rectosigmoid biopsies in collaboration with the Clinical Pharmacology Analytical Laboratory at The Johns Hopkins University School of Medicine as previously described ^24,25^.

### PBMC Isolation

Freshly collected human blood was centrifuged at 1200 rpm for 10 min. Plasma was removed and stored at - 80°C until analysis. Remaining blood was diluted 1:1 with PBS (Thermo Fisher Scientific, Waltham, MA) and added slowly to the side of a SepMate PBMC isolation tube (STEMCELL Technologies, Vancouver, Canada). Overlayed blood was then spun at 1200 g for 10 min. After centrifugation, PBS and blood were removed from the top of the SepMate filter to collect the buffy coat layer, containing the PBMC, washed with PBS, and spun at 1200 rpm for 20 min. The resulting pellet was incubated with 1 mL of 1X Red Blood Cell (RBC) Lysis Buffer, containing 0.2 g NH_4_Cl, 8.4 g NaHCO_3_, 3.7 g EDTA disodium salt in 1 L dH_2_O (Thermo Fisher Scientific, Waltham, MA) for 10 min, washed with PBS, and spun at 1200 rpm for 10 min. The supernatant was aspirated, and PBMC were counted.

PBMC from each participant were lysed in 1X proteomics lysis buffer containing 5% sodium dodecyl sulfate (SDS) (Invitrogen, Waltham, MA) and 50 mM triethylammonium bicarbonate (Sigma-Aldrich, St. Louis, MO) or 70% LC-MS methanol for TFV quantification. TFV-DP was quantified in PBMC lysate from Cohort B in collaboration with the Clinical Pharmacology Analytic Laboratory at The Johns Hopkins University School of Medicine as previously described ^24–27^. The assay lower limits of quantitation was 5 fmol of intracellular TFV- DP/sample. Data were normalized to cell count and represented as fmol/10^6^ cells.

### PBMC TFV Activation Assay

Following isolation, human PBMC from Control Cohort A were cultured in Nunc Bioreactor centrifuge tubes (Thermo Fisher Scientific, Waltham, MA) with PBMC media containing RPMI 1640 medium (Thermo Fisher Scientific, Waltham, MA), 10% FBS (Thermo Fisher Scientific, Waltham, MA), and 1% FBS (Thermo Fisher Scientific, Waltham, MA). PBMC were exposed to 100 μM TFV (Gilead Sciences, Foster City, California) for 4 h at 37°C, 5% CO_2_. TFV incubation was limited to 4 h to minimize PBMC adherence and differentiation. Following incubation, the entirety of the cell culture was removed and set aside. Adhered PBMC were then trypsinized at 37°C, 5% CO_2_ for 20 min and subsequently deactivated with PBMC media. Trypsinized PBMC were collected and added into the cell culture that was set aside following incubation. PBMC were spun at 1200 RPM for 5 minutes, lysed with 70% LC-MS methanol, and frozen at -80°C until analysis. TFV-DP was quantified in PBMC lysate from Control Cohort A in collaboration with the Clinical Pharmacology Analytical Laboratory at The Johns Hopkins University School of Medicine as previously described ^24,25^.

### LC-MS Proteomics

Rectosigmoid tissues were homogenized using a BeadBug bead mill (Benchmark Scientific, Sayreville, NJ) in 1× proteomics lysis buffer containing 5% sodium dodecyl sulfate (SDS) (Invitrogen, Waltham, MA) and 50 mM triethylammonium bicarbonate (Sigma-Aldrich, St. Louis, MO). Tissue lysates were centrifuged at 13,000 x g at 4°C for 20 min to allow proteins to precipitate and pellet. PBMC were lysed in 1x proteomics lysis buffer containing 5% sodium dodecyl sulfate (SDS) (Invitrogen, Waltham, MA) and 50 mM triethylammonium bicarbonate (Sigma-Aldrich, St. Louis, MO). Tissue lysates were processed for proteomics by mass spectrometry and analyzed as previously described ^28^ with the exception that proteins were digested into peptides using the S-Trap 96-well plate (ProtiFi, Fairport, NY) according to the manufacturer’s instructions with the exception that alkylation was performed with methyl methanethiosulfonate (MMTS) rather than iodoacetamide . Resulting spectra were uploaded to Spectronaut 18.7 (Biognosys, Cambridge, MA). Peptides were identified and quantified using the directDIA analysis default settings with the proteotypicity filter set to “Only Proteotypic” and a “methylthio” variable modification. For PBMC proteomics data, MS1 protein group quantifications, UniProt numbers, and molecular weights from Spectronaut (Biognosys, Cambridge, MA) were imported into Perseus (Max Planck Institute of Biochemistry, Planegg-Martinsried, Bavaria, Germany) for use of the Proteomic Ruler plug-in, as previously described^29^. The default proteomic ruler plug-in settings were used for the PBMC. The scaling mode and ploidy were set to the default proteomic ruler plug-in settings. Protein concentrations (nM) and copy numbers, as estimated by the Proteomic Ruler, were used for analysis.

### Pathway analysis

To perform pathway analyses, statistically significant (p < 0.05) enzymes of interest were ranked and represented by -log(p-value) calculated by SimpliFi by Protifi (Fairport, NY) derived from our LC-MS proteomics to assess aging and HIV infection using our human PBMC and rectosigmoid tissues. Pathway categories were arranged and calculated manually by summing relevant -log(p-values) and represented using Prism software 11.0 (GraphPad Software, Inc., San Diego, CA)

### Statistical analysis

A minimum of three independent experiments were performed for PBMC LC-MS proteomics each with two LC- MS injection replicates except for the old, HIV negative group which was limited to two independent experiments. Similarly, a minimum of three independent experiments were performed for rectosigmoid LC-MS proteomics, each with three injection replicates. Independent and technical replicates that contained missing values were omitted from statistical and graphical analyses. All data are graphically represented as means ± SD. Statistical analyses were performed using Prism software 11.0 (GraphPad Software, Inc., San Diego, CA) and R statistical software is v4.6.1. A D’Agostino-Pearson normality test was performed to evaluate whether the data fit a Gaussian distribution. When the data were normally distributed, a Brown-Forsythe and Welch ANOVA was performed. When the data were not normally distributed, a Kruskal-Wallis test was performed. When present, the vehicle treatment condition served as the reference group for multiple comparisons analyses in the one-way ANOVA test instead of a two-way test due to the constraints of missing representative samples in a HIV positive, young group. *p≦0.05. **p≦0.01. ***p≦0.001. ****p≦0.0001.

## Results

### Aging and HIV Infection Differentially Remodel PBMC Proteins Involved in ARV Disposition

Although aging has been associated with altered ARV pharmacokinetics, its impact on the tissue- specific expression of proteins that govern intracellular ARV disposition remains incompletely understood. In particular, little is known about how aging and HIV infection influence drug transporters, metabolizing enzymes, activating kinases, phosphatases, and nucleotide homeostasis pathways within colorectal tissue, a major HIV reservoir. To address this gap, we performed LC-MS-based proteomic profiling of PBMC and rectosigmoid tissue from young HIV-seronegative adults, older HIV-seronegative adults, and older virally suppressed PLWH (**Figures 1-2**). These analyses were designed to define the effects of age, HIV status, and sex on molecular determinants of ARV disposition across biologically distinct compartments ^1^. Our study involved participants from two human cohorts (IRB00307426): young (age 18-50 years old) and old (age 65-80 years old) HIV negative participants (Cohort A) as well as old, HIV positive participants (Cohort B). Cohort A enrolled HIV negative adults in two subsets based upon age, including a young (age 18-50) subset and old (age 65-80) subset and Cohort B enrolled HIV positive adults in an old (age 65-80) group. Cohort A contained 10 young, HIV negative (age 18-50) and three old, HIV negative (age 65-80) adults who were not infected with HIV and not taking tenofovir-based PrEP (“Control Cohort”). Cohort demographics, including age, sex, and race are included in **Table 1**. In Cohort B HIV positive adults received a TFV-based regimen (“ART Cohort”), including FTC, TFV disoproxil fumarate (TDF), TFV alafenamide (TAF), or bictegravir (BIC) in fixed dose combinations (Truvada^TM^, Descovy^TM^, or Biktarvy^TM^; Gilead Sciences, Foster City, California).

**Figure 1.**
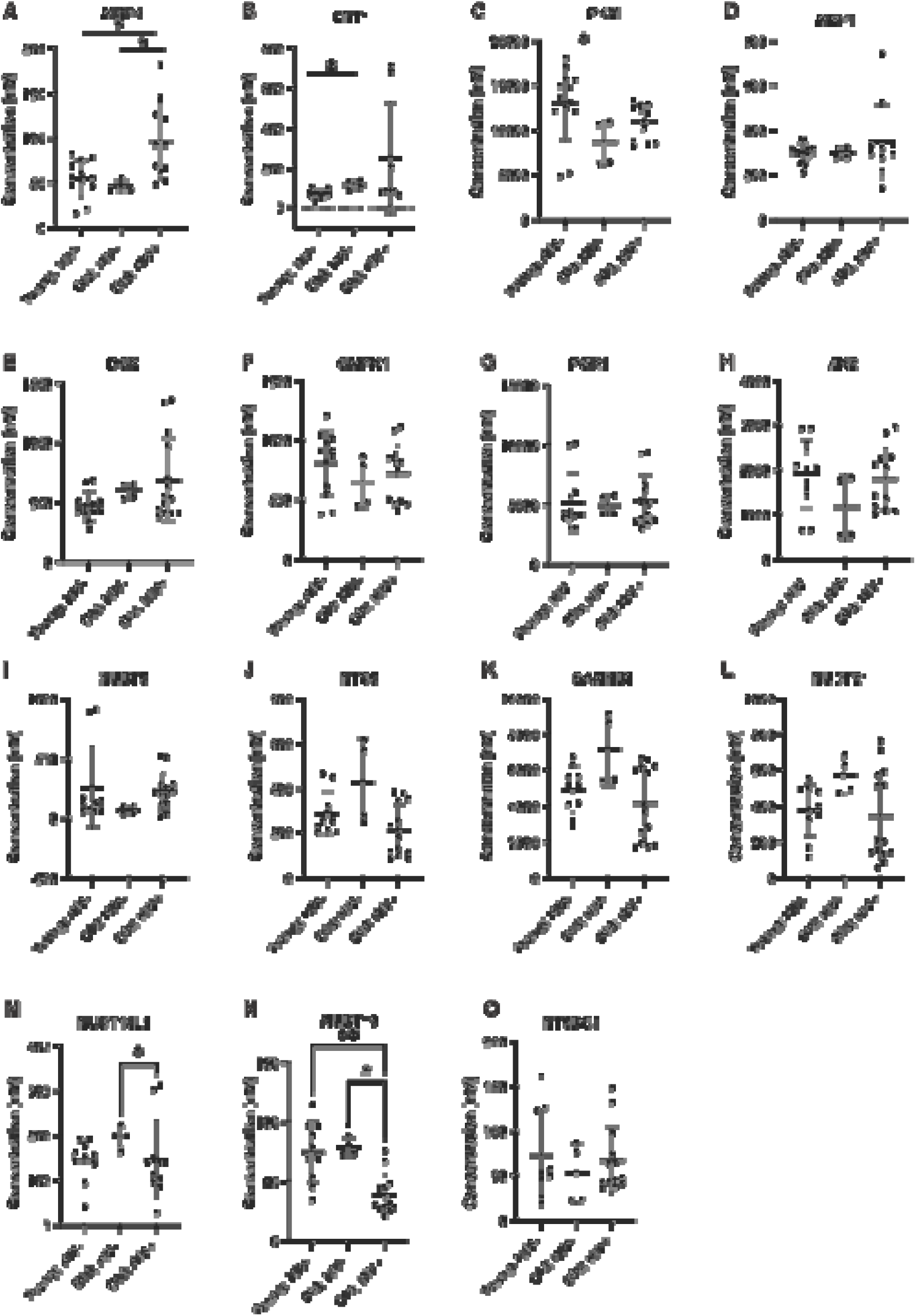
Age and HIV infection affect ARV transporter concentrations in human PBMC. Human PBMC were isolated from young and old, HIV negative participants as well as old, HIV positive participants for measurement of **(A)** MRP4, **(B)** ENT1, (**C)** PKM, **(D)** MRP1, **(E)** DCK, **(F)** CMPK1, **(G)** PGK1, **(H)** AK2, (I) NUDT5, **(J)** NTC5, **(K)** SAMHD1, **(L)** NUDT21, **(M)** NUDT16L1, **(N)** NUDT19, **(O)** NT5DC1 ARV-disposition related transporter and metabolizing enzyme concentrations. All participants in the old, HIV positive cohort were on a TFV-based ART regiment for treatment of HIV. Concentrations were measured by Proteomic Ruler. A minimum of three independent experiments were performed for PBMC LC-MS proteomics each with two LC-MS injection replicates except for the old, HIV negative group which was limited to two independent experiments (represented as individual dots). D’Agostino & Pearson, Anderson-Darling, Shapiro-Wilk, and Kolmogorov-Smirnov test were performed to assess. normality and lognormality. Data represented as mean ± standard deviation. Statistical analysis was performed for normally distributed data using Brown–Forsythe and Welch ANOVA (A, C, F, H) and non-parametric data using Kruskal–Wallis ANOVA (B, D, E, G) by GraphPad software. \**p* < 0.05, \*\**p* < 0.01, \*\*\**p* < 0.001, and \*\*\*\**p* < 0.0001.

**Figure 2.**
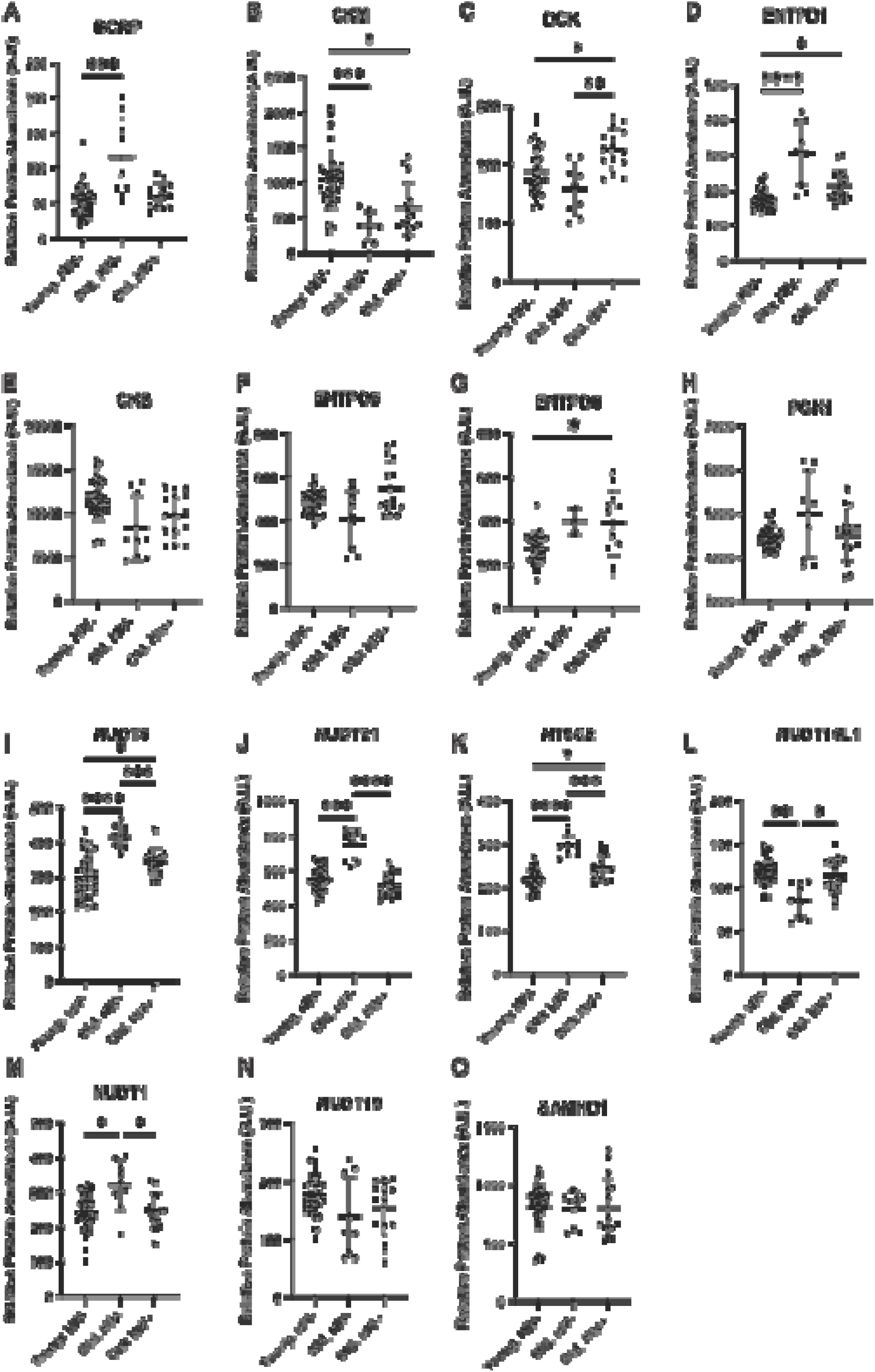
Age and HIV infection impact ARV metabolizing enzyme, transporter, and diphosphohydrolase relative abundances in human rectosigmoid biopsies. Human rectosigmoid biopsies were collected from young and old, HIV negative participants as well as old, HIV positive participants for measurement of ARV related transporters, activators, and metabolizing enzymes including: **(A)** BCRP, **(B)** CKM, **(C)** DCK, **(D)** ENTPD1, **(E)** CKB, **(F)** ENTPD5, **(G)** ENTPD8, **(H)** PGK1, **(I)** NUDT5, **(J)** NUDT21, **(K)** NT5C2, **(L)** NUDT16L1, **(M)** NUDT1, **(N)** NUDT19, **(O)** SAMHD1 were measured by LC-MS proteomics in rectosigmoid biopsies from young and old, HIV negative participants as well as old, HIV positive participants. LC-MS proteomics of rectosigmoid biopsies from young and old, HIV negative participants as well as old, HIV positive participants measured the relative abundance of **(D)** ENTPD1, **(E)** CKB, **(F)** ENTPD5, **(G)** ENTPD8, and **(H)** PGK1 between cohorts. All participants in the old, HIV positive cohort were on a TFV-based ART regiment for treatment of HIV. A minimum of three independent experiments were performed from each cohort (young, HIV negative; old, HIV negative; and old, HIV positive) for colon LC-MS proteomics, each with three injection replicates (represented as individual dots). D’Agostino & Pearson, Anderson-Darling, Shapiro-Wilk, and Kolmogorov-Smirnov test were performed to assess. normality and lognormality. Data represented as mean ± standard deviation. Statistical analysis was performed for normally distributed data using Brown–Forsythe and Welch ANOVA (B, C, E, F, G, I, K, L, M, N) and non- parametric data using Kruskal–Wallis ANOVA (A, D, H, J, O) by GraphPad software. \**p* < 0.05, \*\**p* < 0.01, \*\*\**p* < 0.001, and \*\*\*\**p* < 0.0001.

We observed significant HIV status and aging-dependent differences in ARV-disposition related proteins in PBMC from participants in both cohorts (**Figure 1**), where three proteins had HIV-related effects and two proteins displayed aging-related effects. PBMC from older PLWH had 107.83% higher MRP4 transporter (96.37±42.82 nM; data are denoted as mean ± standard deviation) concentrations as compared to older, HIV negative participants (46.37±5.91 nM; p=0.0089) (**Figure 1A**). In contrast, older PLWH had a 25.34% and 50.00% decrease in nucleotide homeostasis proteins NUDT16L1 (147.3 ± 86.96nM) and NUDT19 (39.59 ± 16.76nM) concentrations, respectively, than older, HIV negative participants (NUDT16L1 [197.3 ± 24.85 nM; p=0.0469] and NUDT19 [79.18 ± 7.21nM, p=0.0196]) (**Figure 1 M-N**). Interestingly, we also observed aging alone, independent of HIV, increased equilibrative nucleoside transporter 1 (ENT1) by 60.130% and decreased PKM by 34.341% in PBMC from older, HIV negative particpants (115.39 ± 20.32 nM ENT1, 8593.47 ± 2531.68 PKM), relative to younger individuals (72.06±19.88 nM, p=0.0322 ENT1, 13,088.03 ± 4,014.84 nM, p = 0.0773 PKM) (**Figure 1 B-C**). In contrast, the other evaluated transporters (MRP1), activating kinases (DCK, CMPK1, PGK1, and AK2), or nucleotide homeostasis proteins (NUDT5, NTC5, SAMHD1, NUDT21, and NT5DC1) remained unchanged (**Figure 1 D,E,F,G,H,I,J,K,L,O**).

### Aging Predominantly Alters ARV Disposition Pathways in Rectosigmoid Tissue

We next evaluated ARV-disposition related proteins in rectosigmoid colon tissue, a major site of HIV acquisition and viral persistence ^30–32^. In contrast to PBMC, aging exerted a substantially greater effect on ARV-disposition related proteins in rectosigmoid tissue, whereas HIV status alone was not associated with significant changes in any evaluated protein (**Figure 2**). Aged participants, irrespective of HIV status, had significantly increased drug transporters (BCRP 115.57% [307.7 ± 91.55 A.U. older, HIV- vs. 173.75 ± 27.33 A.U. young, HIV-, p < 0.0001]) (**Figure 2A**), phosphatases (ENTPD1 77.14% [307.7 ± 91.55 A.U. older, HIV- vs 173.75 ± 27.33 A.U. young, HIV-, p < 0.0001] and ENTPD8 43.621% [395.33 ± 147.71 A.U. older, HIV+ vs 275.26 ± 14.726 A.U. young, HIV-; p = 0.0311]) (**Figure 2 D,G**), and several nucleotide homeostasis proteins (NUDT5 38.38% [416.8±32.7 A.U. older,HIV- vs 301.2±61.93 A.U. young,HIV-, p < 0.0001], NUDT21 35.96% [752.0±83.12 A.U. older,HIV- vs 553.1±72.29 A.U. young,HIV-, p=0.0004], NT5C2 35.67% [298.6±23.32 A.U. older,HIV- vs 220.1±23.37 A.U. young,HIV-, p < 0.0001], and NUDT1 [322.0±72.96 A.U. older,HIV- vs 242.5±54.92 A.U. young,HIV- p=0.0340]) (**Figure 2 I-K,M**) while decreasing the activating enzyme (CKM 39.52% [634.16 ± 367.69 A.U. older, HIV+ vs 1048.46 ± 422.42 A.U. young, HIV-, p = 0.0145] and increasing the activating enzyme DCK 19.82% [226.7±34.59 A.U. older, HIV+ vs 189.2±42.19 A.U. young, HIV-, p=0.0184]) (**Figure 2B-C**). In contrast, aging in the HIV-negative participants decreased the nucleotide homeostasis protein NUDT16L1 26.57% (87.16±14.86 A.U. older, HIV- vs 118.7±14.86 A.U. young, HIV-, p=0.0031) (**Figure 2 L**). Interestingly, while some age-associated differences were maintained in rectosigmoid tissue obtained from older PLWH (CKM and ENTPD8) (**Figure 2 B,G**), others were attenuated or reversed in the setting of HIV infection, including BCRP, DCK, ENTPD1, NUDT5, NUDT21, NT5C2, NUDT16L1, and NUDT1(**Figure 2 A,C,D, I-K,L,M**). In contrast, the activating kinase CKB and the nucleotide homeostasis proteins (NUDT19 and SAMHD1) remained unchanged in rectosigmoid tissue across all groups (**Figure 2 E,N,O**). Together, these findings indicate that aging is the primary factor associated with altered expression of proteins governing ARV disposition in rectosigmoid tissue, with HIV infection modulating only a subset of these age-related effects.

### Sex Modifies Age- and HIV-Associated Differences in Rectosigmoid ARV Disposition Proteins

As substantial hormonal changes occur across the lifespan that may influence ARV disposition, we next evaluated whether sex modified the age- and HIV-associated differences observed in rectosigmoid tissue (**Figure 2**). We identified several sex-dependent effects among proteins involved in ARV activation, metabolism, and transport (**Figure 3**). Notably, younger HIV-negative women exhibited significantly higher concentrations of the activating kinases CKB and CKM, both of which progressively declined with aging compared to male participants (CKB: 26.61% decrease, 13877±1311 A.U. female, young, HIV- vs 10184±1689 A.U. male,young,HIV-, p=0.0005; CKM: 60.92% decrease, 1219±491.1 A.U. female, young, HIV- vs 476.4±143.4 A.U., female, old, HIV- p=0.0119) (**Figure 3 A,B**). HIV-status only had a robust impact on the activating enzyme DCK. Older HIV-negative women had lower concentrations of the activating kinase DCK compared with older men living with HIV (30.61% decrease, 157.3±42.0 A.U. vs 226.7±34.59 A.U., p=0.0344) (**Figure 3E**). Interestingly, age-associated increases in the phosphatase ENTPD1 and the transporter BCRP were most pronounced in women, indicating a sex-dependent amplification of aging-related effects on proteins regulating intracellular ARV disposition (**Figure 3 F,G**). Interestingly, the activating kinase, AK2, remained unchanged across all sex, age, and HIV-status groups (**Figure 3D**). Together, these findings suggest that biological sex modifies several age-associated changes in rectosigmoid proteins involved in ARV activation, transport, and metabolism, highlighting sex as an additional contributor to variability in tissue-specific ARV disposition.

**Figure 3.**
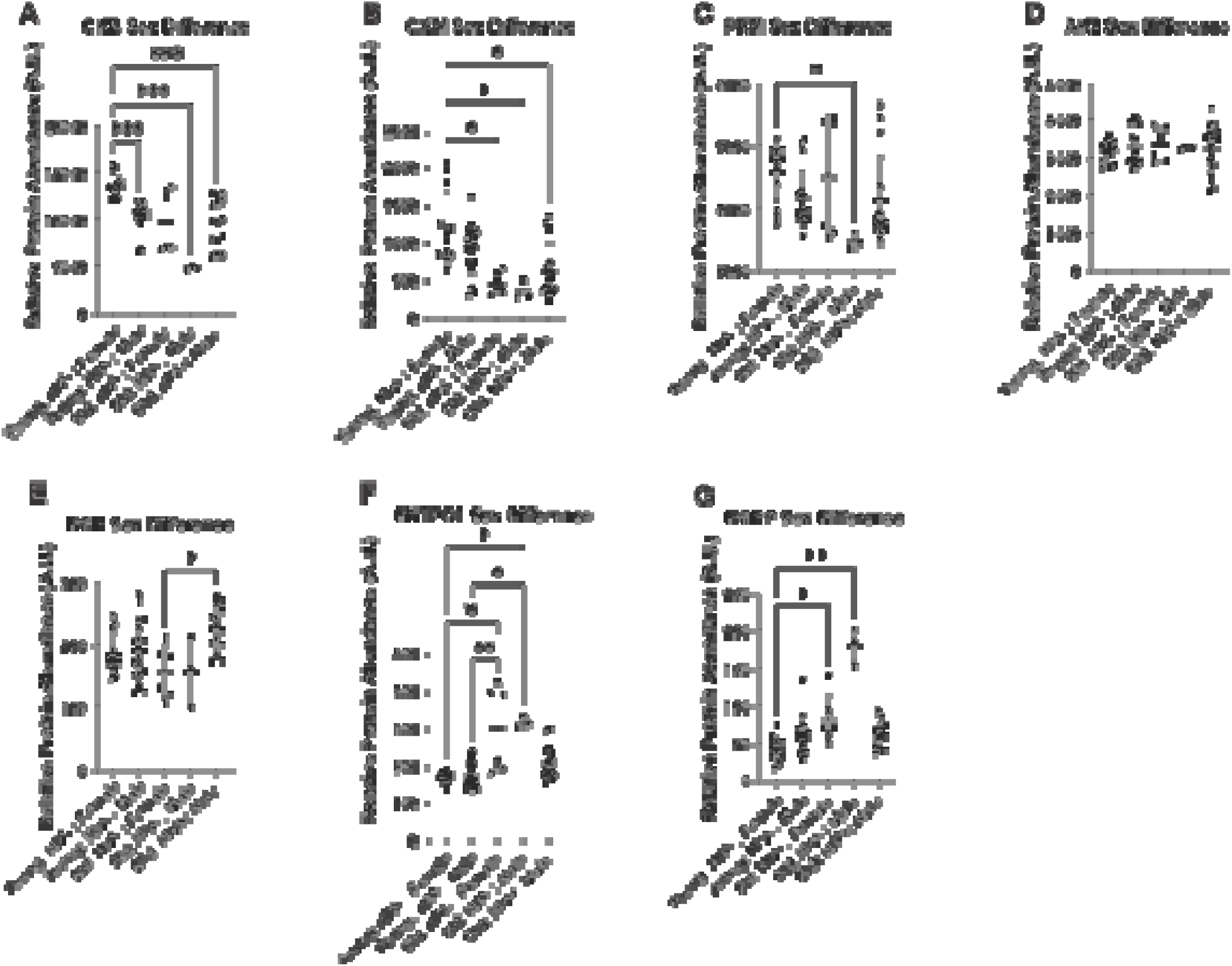
ARV-metabolizing enzymes abundance shifts at the intersection of sex, aging, and HIV status. Human rectosigmoid colorectal tissues were isolated from males and females of young and old, HIV negative participants as well as old, HIV positive participants for measurement of **(A)** CKB, **(B)** CKM, **(C)** PKM, **(D)** AK2, **(E)** DCK, **(F)** ENTPD1, **(G)** BCRP concentrations. All participants in the old, HIV positive cohort were on a TFV-based ART regimen for treatment of HIV. Concentrations were measured by Proteomic Ruler. A minimum of three independent experiments were performed for rectosigmoid LC-MS proteomics each with two LC-MS injection replicates except for the old, HIV negative group which was limited to two independent experiments (represented as individual dots). D’Agostino & Pearson, Anderson-Darling, Shapiro-Wilk, and Kolmogorov- Smirnov test were performed to assess. normality and lognormality. Data represented as mean ± standard deviation. Statistical analysis was performed using Kruskal–Wallis ANOVA by GraphPad for nonparametric data software. *p < 0.05, **p < 0.01, ***p < 0.001, and **** p < 0.0001.

### *In vitro* TFV Activation Assay Does Not Mirror *Ex vivo* Proteomic Remodeling Across Age and HIV Status

Next, we evaluated the functional consequences of the age and HIV infection-driven changes in ARV-disposition related proteins by performing a TFV activation assay in PBMC isolated from young and old HIV negative participants (**Figure 4A**). In this assay, we evaluated TFV activation capacity by exposing isolated PBMC *in vitro* to TFV for 4 h and measuring the resulting TFV-DP formation by LC-MS (**Figure 4A**), according to our established method ^26^. We then compared our calculated TFV-DP formation to clinical TFV-DP concentrations from old, HIV positive participants (**Figure 4A**). In PBMC, we did not observe significant differences in metabolism between young and old, HIV negative participants nor old, HIV positive participants, as indicated by comparable TFV-DP concentrations (**Figure 4A**). Next, we confirmed TFV metabolism in rectosigmoid biopsies isolated from participants on TFV-based regimens, including old, HIV positive participants (Cohort B) by measuring TFV-DP concentrations using LC-MS (**Figure 4B,C**). TFV and TFV-DP concentrations were measured in all five old, HIV positive participants; however, significant interindividual variability was present (**Figure 4B,C**). The source of this variability was not due to sex-based differences as all participants were male. Differences in TFV metabolism due to age were not evaluated in rectosigmoid biopsies from young and old, HIV negative participants as they were not on any ART. Nevertheless, despite low sample number for our NRTI pharmacokinetics, our proteomics data demonstrate protein and site-specific effects of age and HIV infection which may impact regulation of and activity of ARV transport, metabolizing, and eliminating proteins.

**Figure 4.**
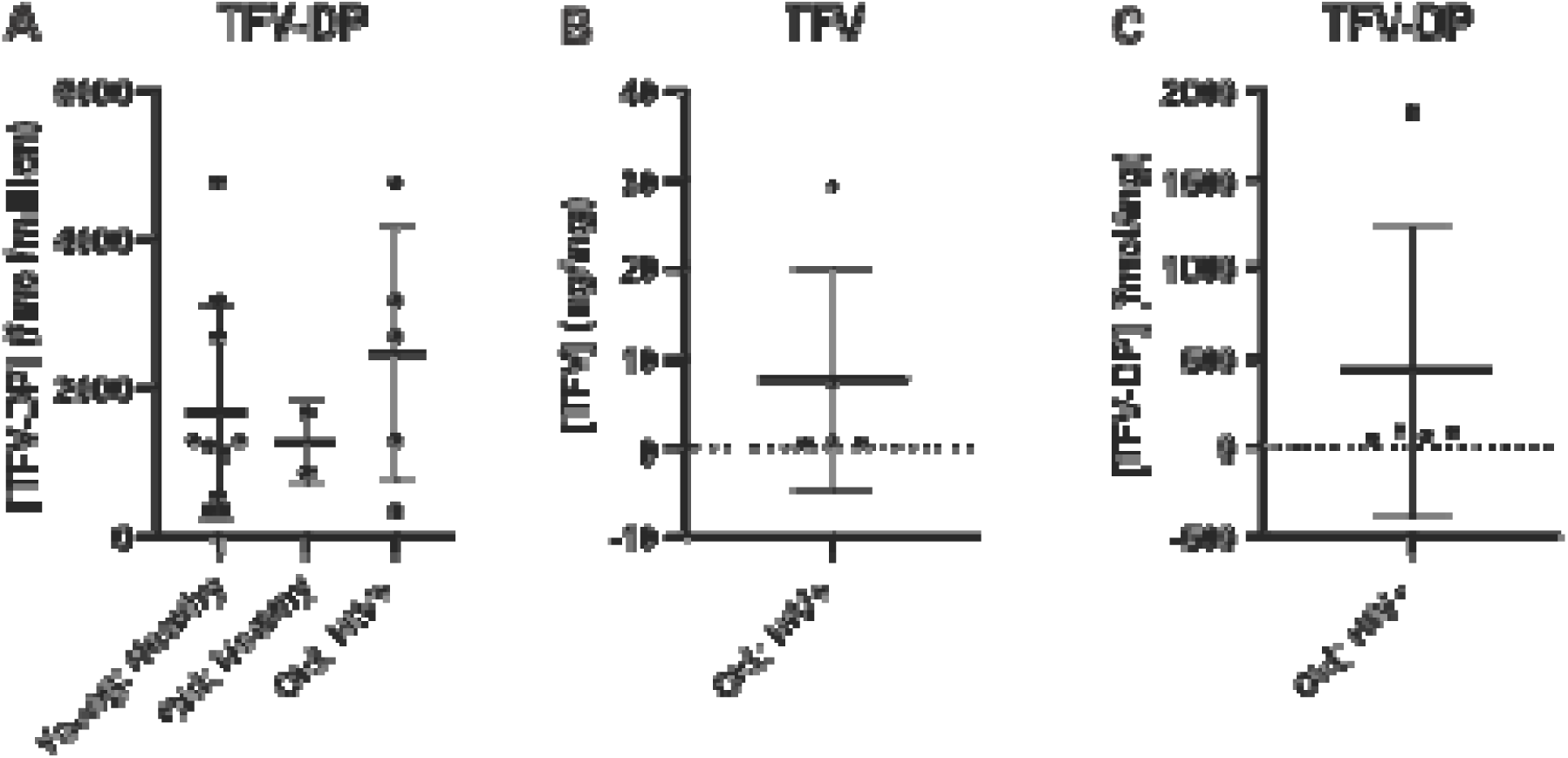
Concentrations of TFV and TFV-DP in human rectosigmoid biopsies and PBMC. TFV-DP concentrations **(A)** were formed from in vitro TFV activation in human PBMC isolated from young and old, HIV negative participants and compared to clinical concentrations of TFV-DP in PBMC from old, HIV positive participants. Isolated HIV negative PBMC were incubated with 100 μM TFV for 4 h at 37°C, 5% CO2 to measure TFV activation to TFV-DP by LC-MS. Concentrations of **(B)** TFV and **(C)** TFV-DP were measured in human rectosigmoid biopsies by LC-MS from old, HIV positive participants on a TFV-based ART regimen. Limits of quantification were concentrations less than 5 fmol/sample (TFV-DP) and 0.05 ng/sample (TFV). A minimum of three independent experiments were performed from each cohort (young, HIV negative; old, HIV negative; and old, HIV positive) for TFV and TFV-DP measurements in rectosigmoid biopsies (represented as individual dots). D’Agostino & Pearson, Anderson-Darling, Shapiro-Wilk, and Kolmogorov-Smirnov test were performed to assess. normality and lognormality. Data represented as mean ± standard deviation. Statistical analysis for TFV-DP (A) was performed using Kruskal–Wallis ANOVA for non by GraphPad software. \**p* < 0.05, \*\**p* < 0.01, \*\*\**p* < 0.001, and \*\*\*\**p* < 0.0001.

### Pathway Analysis Identifies Broad Tissue-Specific Remodeling Associated with Aging and HIV Infection

ARV-disposition related proteins function within larger interconnected cellular networks and it is likely that changes in their abundance may reflect broader alterations in tissue physiology. To provide additional mechanistic context, we performed pathway analyses of the full proteomic dataset to determine how aging and HIV infection influence pathways relevant to ARV disposition, immune function, disease pathology, and cellular metabolism. Statistically significant pathways (p<0.05) were identified using SimpliFi and ranked by -log(p- value) derived from LC-MS proteomic data obtained from PBMC and rectosigmoid tissues. Across both tissues, the most significantly altered pathways clustered into four major categories: 1) ARV metabolism and transport, 2) disease-associated processes, 3) immune response and signaling, and 4) cellular energy metabolism (**Figure 5,6**,**7**,**8**). We determined that resulting altered pathways across HIV status and aging groups in both PBMC and rectosigmoid tissue were relevant to 1) drug metabolism and transport, 2) pathological disease responses 3) immune response and signaling, and 4) cellular and organismal energy metabolism (**Figures 5-8**).

**Figure 5.**
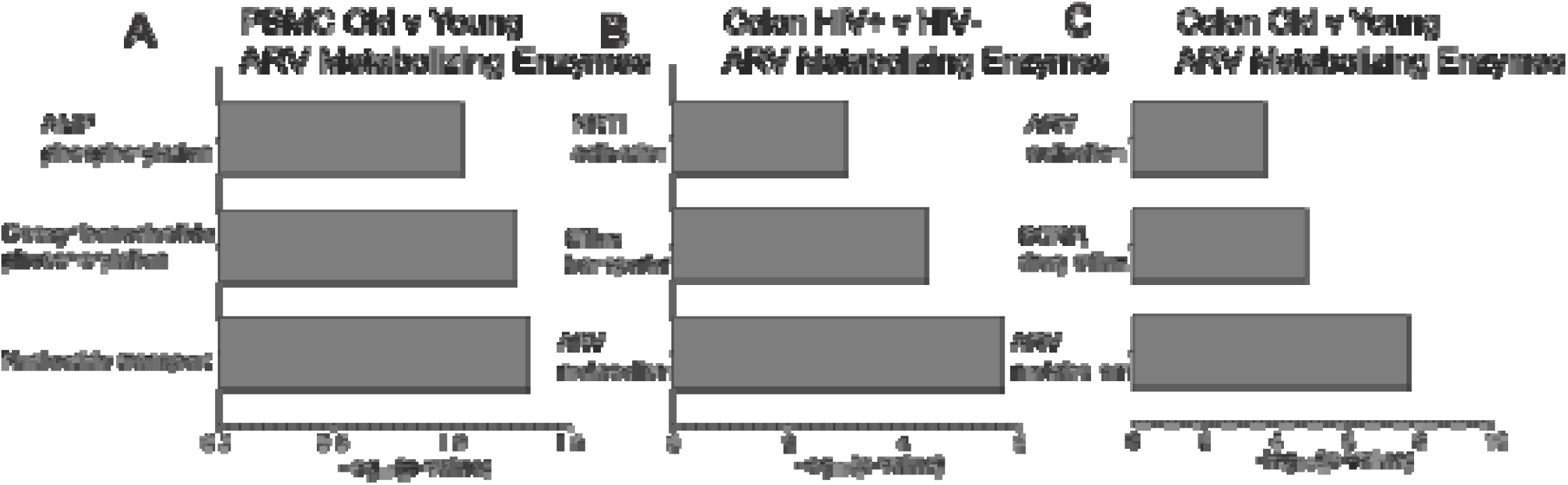
ARV metabolizing enzyme pathways are altered due to aging in PBMC and HIV infection and aging in rectosigmoid tissue. Proteomics were measured and pathway analysis conducted using human PBMC and rectosigmoid biopsies from young, HIV negative and old, HIV negative participants as well as old, HIV positive participants. All participants in the old, HIV positive, old cohort were on a TFV-based ART regimen for treatment of HIV. Pathway analyses for drug metabolism related enzymes were represented for **(A)** old, HIV negative vs. young, HIV negative PBMC, **(B)** old, HIV positive vs. old, HIV negative rectosigmoid tissue, and **(C)** old, HIV negative vs young, HIV negative rectosigmoid tissue. Data represented as -log_10_(p-value) using GraphPad software. P-value and proprietary proteomics analyses generated by Simplifi by Protifi. Data represented as -log_10_(p-value) using GraphPad software. P-value and proprietary proteomics analyses generated by Simplifi by Protifi.

**Figure 6.**
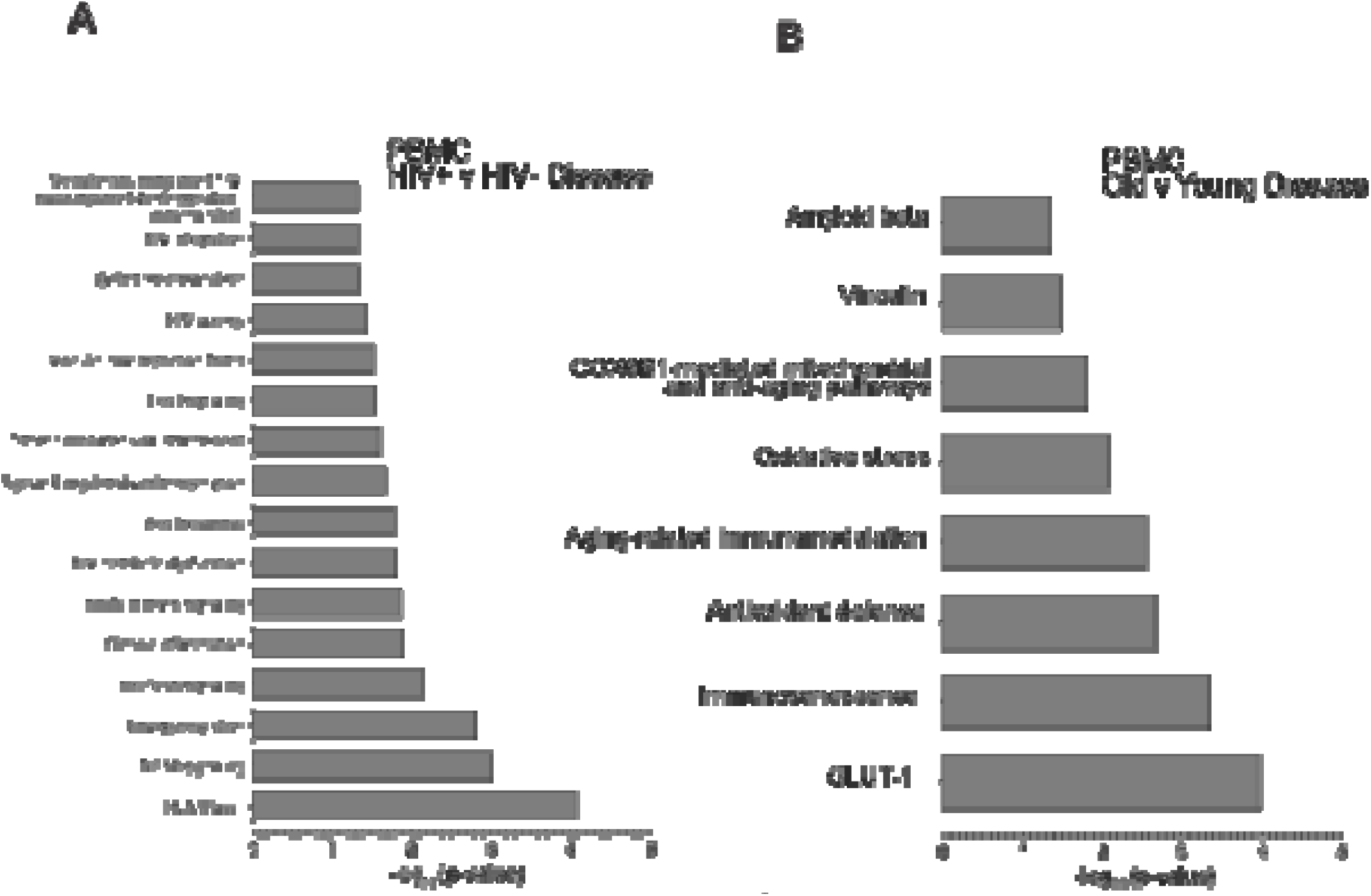
Disease-related pathways are impacted by both HIV and aging in PBMC. Proteomics were measured and pathway analysis conducted using human PBMC and rectosigmoid biopsies from young, HIV negative and old, HIV negative participants as well as old, HIV positive participants. All participants in the old, HIV positive cohort were on a TFV-based ART regimen for treatment of HIV. Pathway analyses for disease related enzymes were represented for **(A)** old, HIV positive vs. old, HIV negative PBMC and **(B)** old, HIV negative vs. young, HIV negative PBMC. Data represented as -log_10_(p-value) using GraphPad software. P- value and proprietary proteomics analyses generated by Simplifi by Protifi.

**Figure 7.**
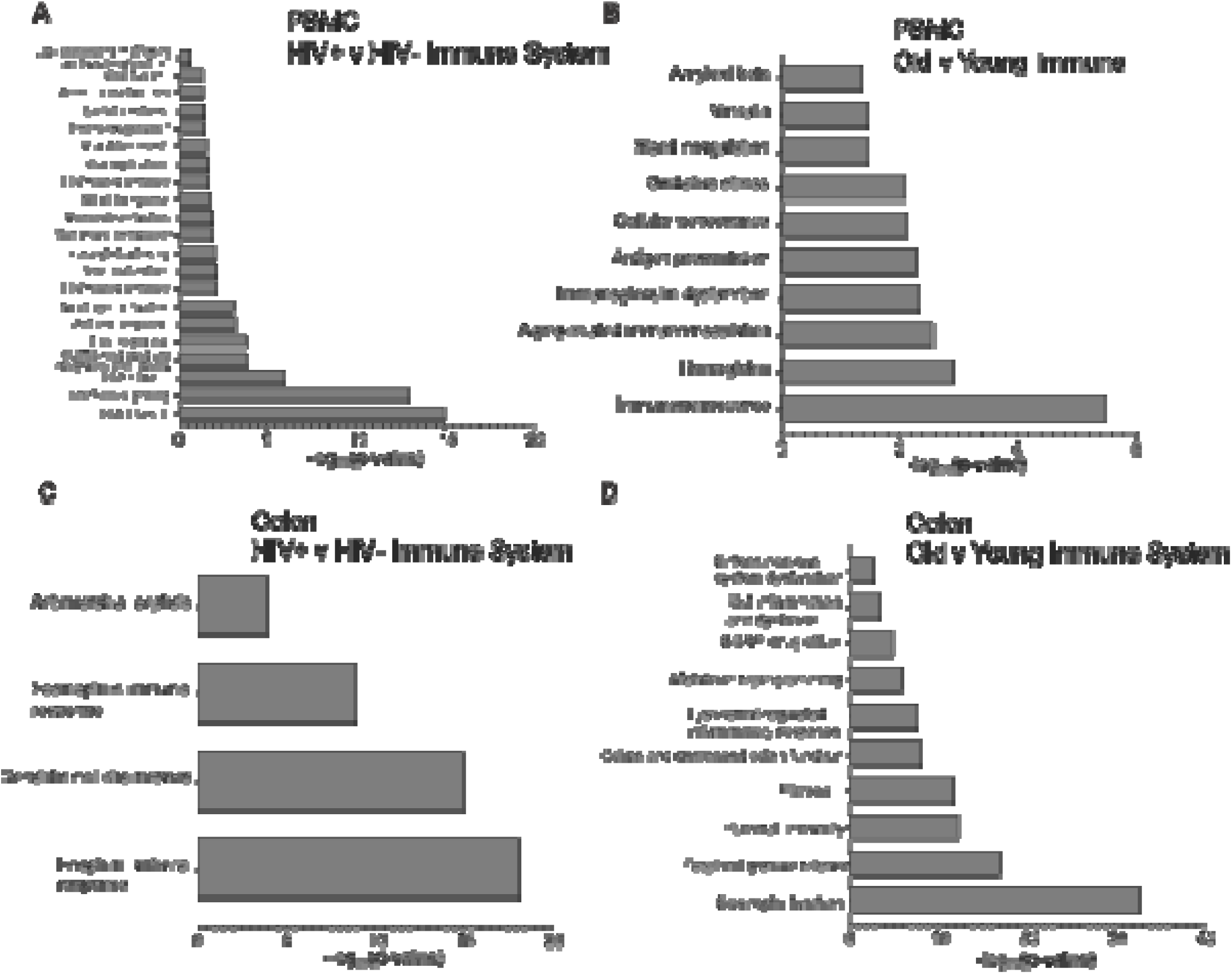
Molecular pathways related to the immune system are impacted broadly in PBMC and rectosigmoid tissue due to HIV infection and aging. Proteomics were measured and pathway analysis conducted using human PBMC and rectosigmoid biopsies from young, HIV negative and old, HIV negative participants as well as old, HIV positive participants. All participants in the old, HIV positive cohort were on a TFV-based ART regimen for treatment of HIV. Pathway analyses for disease related enzymes were represented for **(A)** old, HIV positive vs. old, HIV negative PBMC, **(B)** old, HIV negative vs. young, HIV negative PBMC, **(C)** old, HIV positive vs. old, HIV negative rectosigmoid tissue, and **(D)** old, HIV negative vs. young, HIV negative rectosigmoid tissue. Data represented as -log_10_(p-value) using GraphPad software. P-value and proprietary proteomics analyses generated by Simplifi by Protifi.

**Figure 8.**
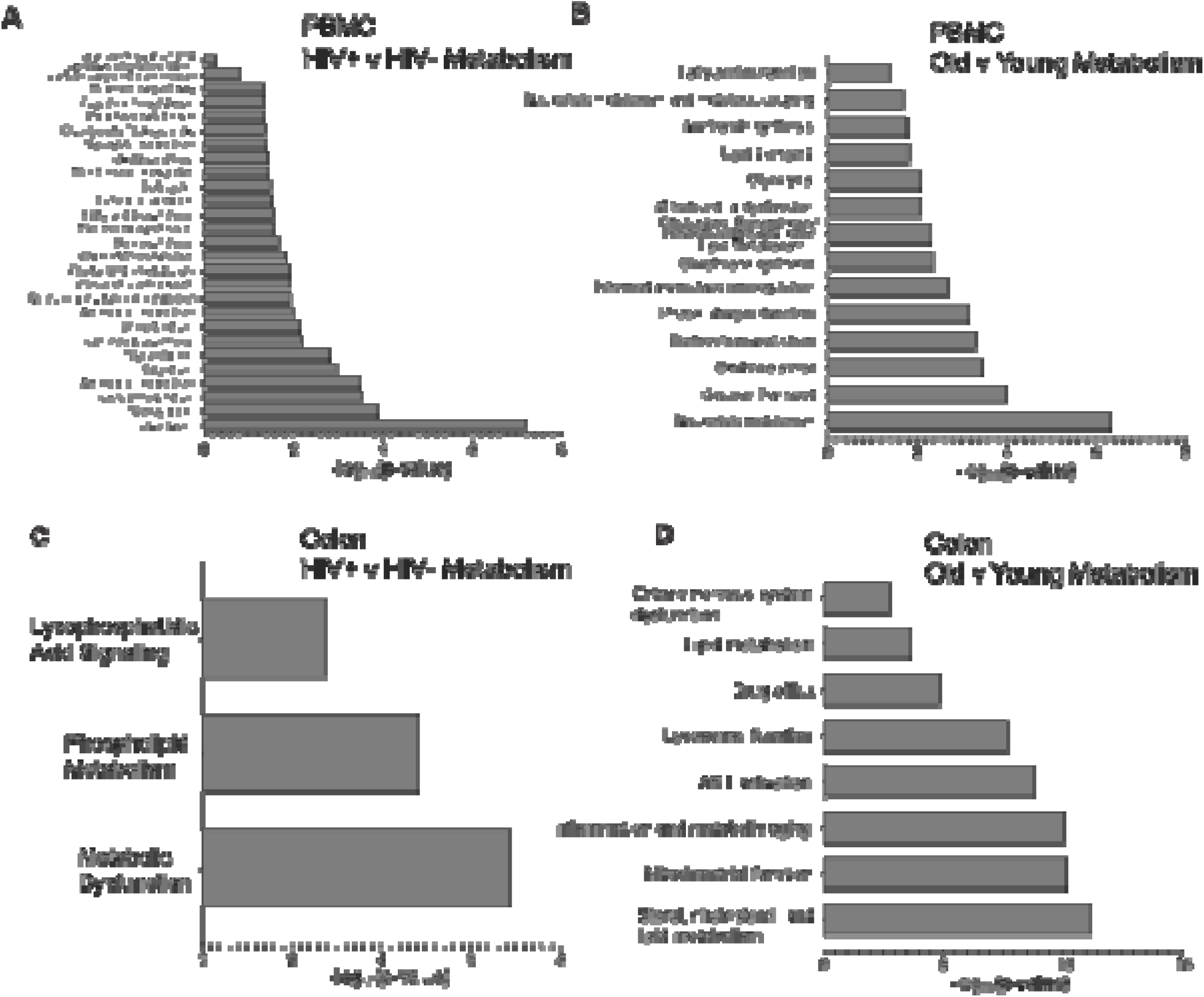
Distinct metabolic pathways are altered due to HIV infection and aging in PBMC and rectosigmoid tissue. Proteomics were measured and pathway analysis conducted using human PBMC and rectosigmoid biopsies from young and old, HIV negative participants as well as old, HIV positive participants. All participants in the old, HIV positive cohort were on a TFV-based ART regimen for treatment of HIV. Pathway analyses for disease related enzymes were represented for **(A)** old, HIV positive vs. old, HIV positive PBMC, **(B)** old, HIV negative vs. young, HIV negative PBMC, **(C)** old, HIV positive vs. old, HIV negative rectosigmoid tissue, and **(D)** old, HIV negative vs. young, HIV negative rectosigmoid tissue. Data represented as -log_10_(p-value) using GraphPad software. P-value and proprietary proteomics analyses generated by Simplifi by Protifi.

Consistent with our protein-specific analyses, pathways involved in ARV activation, transport, clearance, and nucleotide metabolism were significantly altered in both PBMC and rectosigmoid tissue (**Figure 5**). However, the relative contributions of aging and HIV infection differed substantially between compartments. Aging altered multiple pathways associated with ARV metabolism in both PBMC and rectosigmoid tissue (**Figure 5A,C**), whereas HIV infection primarily affected pathways involved in ARV transport, activation, clearance, and metabolism in rectosigmoid tissue from older participants (**Figure 5B**). Together, these findings suggest that both aging and HIV infection influence molecular pathways relevant to ARV disposition, although the magnitude and tissue-specific patterns of these effects differ.

We next evaluated disease-related pathways that may contribute to, or result from, the observed changes in ARV-disposition related proteins (**Figure 6**). In PBMC, HIV infection produced the broadest effects, significantly altering pathways associated with chronic inflammation, HIV metabolic dysfunction, viral latency, and HIV integration (**Figure 6A**). Aging also modified disease-related pathways in PBMC, including immunosenescence, antioxidant defense, oxidative stress, and age-associated immunomodulation, although these effects were less extensive than those associated with HIV infection (**Figure 6B**). In contrast, relatively few disease-associated pathways were altered by either aging or HIV infection in rectosigmoid tissue, suggesting that disease-related proteomic remodeling is more pronounced in circulating immune cells than within the colorectal mucosa.

Because both aging and HIV are known to alter immune function, we next examined immune response and signaling pathways (**Figure 7**). Similar to the disease-related analyses, PBMC exhibited the broadest immune pathway alterations. HIV infection significantly modified pathways associated with interferon signaling, T-cell signaling, antiviral responses, leukocyte trafficking, monocyte activation, and B-cell responses (**Figure 7A**). Aging also impacted numerous immune-related pathways in PBMC, including immunosenescence, age- related immunomodulation, antigen presentation, cellular senescence, and oxidative stress responses (**Figure 7B**). In rectosigmoid tissue, HIV infection induced comparatively modest changes that were primarily associated with eosinophil-mediated antiviral responses, dendritic cell chemotaxis, and antimicrobial peptide pathways (**Figure 7C**). In contrast, aging altered a broader range of immune-related pathways in rectosigmoid tissue, including eosinophil function and granule release, humoral immunity, fibrosis, colitis-associated responses, lysosome-regulated inflammation, metabolic reprogramming, gut inflammation and dysbiosis, enteric nervous system dysfunction, and BCRP-mediated drug efflux (**Figure 7D**). These findings suggest that aging may exert a greater influence on immune-related processes in colorectal tissue than HIV infection itself.

Finally, we examined pathways involved in cellular energy metabolism (**Figure 8**). Similar to the patterns observed for disease and immune pathways, HIV infection produced the most extensive metabolic remodeling in PBMC. Significantly altered pathways included carbohydrate metabolism, tricarboxylic acid cycle activity, amino acid metabolism, electron transport chain function, oxidative stress regulation, prostaglandin metabolism, carnitine biosynthesis, glycosylation, fatty acid biosynthesis, and fatty acid oxidation (**Figure 8A**). Aging more prominently affected pathways associated with nucleotide synthesis and metabolism, metabolic coupling, and cholesterol and isoprenoid biosynthesis in PBMC (**Figure 8B**). In rectosigmoid tissue, HIV infection exerted more limited effects that were largely restricted to pathways associated with metabolic dysfunction, phospholipid metabolism, and lysophosphatidic acid signaling (**Figure 8C**). In contrast, aging substantially altered pathways involved in enteric nervous system dysfunction, lipid metabolism, mitochondrial function, sterol metabolism, inflammation-associated metabolic remodeling, drug efflux, and ARV activation in rectosigmoid tissue (**Figure 8D**). Collectively, these findings demonstrate that aging and HIV infection are associated with extensive but tissue-specific proteomic remodeling and support the hypothesis that broader alterations in immune, metabolic, and drug disposition pathways may contribute to the age- and HIV- associated differences observed in ARV-disposition related proteins. As older adults remain underrepresented in many ART efficacy studies, these findings provide a mechanistic rationale for future clinical investigations examining whether age-associated biological changes influence ARV disposition and treatment outcomes in aging populations ^7,8^.

## Discussion

Transporters, metabolizing enzymes, activating kinases, phosphatases, and nucleotide homeostasis proteins collectively regulate intracellular ARV disposition by controlling drug uptake, activation, catabolism, and elimination. Aging is known to alter the expression and activity of many of these proteins, contributing to a dynamic pharmacologic environment that may be further modified by HIV infection, comorbid disease, and polypharmacy ^13^. Previous studies reported age-dependent changes in the abundance or function of ARV- relevant solute carrier (SLC) and MRP transporters, cytochrome P450 enzymes, UDP- glucuronosyltransferases (UGTs), and the TFV-activating kinase CKM ^16,23,33–39^. However, most investigations focused on blood-derived samples or systemic pharmacokinetics, leaving important questions regarding age- associated changes within HIV reservoir tissues unresolved. Here, we demonstrate that aging, HIV status, and biological sex influence multiple determinants of ARV disposition, with the most pronounced effects occurring in rectosigmoid tissue rather than PBMC.

Overall, we determined that aging exerted a substantially greater influence on ARV-disposition related proteins in rectosigmoid tissue than HIV status alone. While relatively few ARV-disposition related proteins were altered in PBMC, aging modified multiple protein classes within rectosigmoid tissue, including transporters (BCRP), activating enzymes (CKM and DCK), phosphatases (ENTPD1 and ENTPD8), and nucleotide homeostasis proteins (NUDT5, NUDT21, NT5C2, NUDT16L1, and NUDT1). These observations are particularly relevant because the gastrointestinal tract serves as both a major site of HIV acquisition and a persistent viral reservoir. Notably, HIV infection alone had limited effects on ARV-disposition related protein abundance in rectosigmoid tissue, suggesting that biologic aging may have a greater influence than HIV status itself on molecular pathways governing local ARV disposition in this compartment.

Several age-associated changes identified in this study are supported by previous observations, whereas others appear to represent novel findings. CKM abundance was reduced in rectosigmoid tissue from older participants, extending prior reports that creatine kinase activity declines with age in serum and skeletal muscle ^22,40^. This observation may be pharmacologically important because CKM contributes to conversion of TFV-MP to its active metabolite, TFV-DP, in colorectal tissue ^41^. Conversely, we observed increased abundance of BCRP and ENTPD1 with aging. Prior studies reported lower hepatic BCRP expression in older adults ^20^, indicating that age-associated regulation of BCRP differs across tissues. Such tissue-specific effects are biologically plausible given known differences in transporter expression and function among distinct anatomic compartments ^17^. However, the functional implications of altered ENTPD remain less clear. ENTPD family members regulate nucleotide phosphate turnover and contribute to broader nucleotide homeostasis pathways that influence nucleotide availability within tissues ^42–44^. Because intracellular phosphohydrolases can affect the persistence of phosphorylated nucleotide analogs, age-associated differences in ENTPD expression warrant further investigation in the context of ARV pharmacology. Collectively, these findings support the concept that aging regulates ARV disposition in a protein- and tissue-specific manner rather than through a uniform mechanism.

One of the most unexpected findings was the prominent effect of aging on proteins involved in nucleotide homeostasis. In addition to changes in transporters and activating enzymes, aging significantly altered several nucleotidases and NUDIX hydrolases, including NUDT5, NUDT21, NT5C2, NUDT16L1, and NUDT1. Although these proteins have received comparatively little attention within HIV pharmacology, they collectively regulate endogenous nucleotide pools and the metabolism of nucleotide analogs. This finding is potentially important because intracellular nucleotide abundance influences competition between endogenous nucleotides and active NRTI metabolites at the level of reverse transcriptase ^45,46^. Consistent with these observations, our pathway analyses independently identified age-associated alterations in nucleotide metabolism pathways. Together, these findings suggest that remodeling of nucleotide homeostasis pathways may represent an underappreciated mechanism contributing to variability in NRTI pharmacodynamics during aging and highlight an area requiring further mechanistic investigation.

Our data additionally suggest that biological sex modifies age-associated changes in proteins involved in ARV disposition. Sex-dependent differences were most evident in rectosigmoid tissue, where CKB, CKM, DCK, ENTPD1, and BCRP displayed distinct patterns across age and HIV-status groups. Younger HIV- negative women generally exhibited higher abundance of several activating enzymes, whereas age-associated increases in ENTPD1 and BCRP were amplified in men. Although the mechanisms underlying these observations remain unclear, age-associated hormonal changes may contribute to differential regulation of proteins involved in ARV activation, transport, and nucleotide metabolism. While these findings require validation in larger cohorts, they suggest that biological sex may influence how aging affects tissue-specific ARV disposition and support inclusion of sex-stratified analyses in future pharmacologic investigations.

An important consideration is the magnitude of the observed proteomic differences and whether they are likely to be clinically meaningful. Proteomic analyses alone cannot determine whether a given change in protein abundance is sufficient to alter drug exposure or ARV efficacy because downstream effects depend on protein activity, substrate affinity, compensatory metabolic pathways, cell-specific expression patterns, and local drug concentrations. Nevertheless, several of our observed changes were substantial. For example, there was a more than a two-fold increase in BCRP abundance within rectosigmoid tissue in the context of aging. Given the broad substrate profile of BCRP and its relevance to multiple ARV classes, the magnitude of this age-associated increase suggests that it may warrant particular attention in future functional studies. Ultimately, determining whether these molecular changes translate into clinically meaningful differences in ARV exposure or response will require dedicated pharmacokinetic and pharmacodynamic investigations. Moreover, our observed proteomic changes were concentrated within pathways governing NRTI transport, activation, dephosphorylation, and nucleotide homeostasis, particularly for TFV- and FTC-related pathways. With the exception of BCRP, which has substrates spanning multiple ARV classes, the findings reported here are not currently expected to have broad implications for non-NRTI ARV.

To place these protein-specific findings into a broader biological context, we performed pathway analyses of the full proteomic dataset. These analyses demonstrated that the observed changes in ARV- disposition related proteins occurred within a larger framework of tissue remodeling involving pathways related to drug metabolism, disease pathology, immune signaling, and cellular metabolism. Interestingly, HIV infection and aging exhibited compartment-specific effects. HIV infection exerted the strongest influence on PBMC, where pathways associated with chronic inflammation, immune activation, viral latency, HIV integration, and cellular metabolism were prominently altered. In contrast, aging produced a broader impact on rectosigmoid tissue, influencing pathways related to gut inflammation, fibrosis, dysbiosis, enteric nervous system function, mitochondrial biology, lipid metabolism, and drug disposition ^30,47,48^. These findings suggest that age- associated alterations in ARV-disposition related proteins occur within a broader landscape of biological remodeling that may influence local drug activity within HIV reservoir tissues.

Because we observed changes in proteins involved in ARV activation and transport, we evaluated TFV activation to TFV-DP in PBMC and rectosigmoid tissue to determine whether these proteomic changes were associated with detectable functional differences. Surprisingly, we observed no detectable differences in TFV activation within PBMC attributable to aging or HIV status despite differences in proteins implicated in TFV metabolism. These findings suggest that modest alterations in individual proteins may not necessarily translate into measurable differences in TFV activation *ex vivo*, potentially because of redundancy and compensatory mechanisms within nucleotide metabolic networks. We additionally confirmed the presence of TFV and TFV- DP formation within rectosigmoid tissue, consistent with previous reports ^49^. However, larger studies will be required to determine whether the proteomic alterations identified here influence ARV exposure, persistence, or antiviral activity within relevant tissue compartments ^47^.

This study has several important limitations. First, the study was exploratory in nature and enrolled the maximum number of participants available during the study period rather than being powered to detect predefined clinical endpoints. Consequently, the relatively small sample size limits statistical power and increases the possibility of false-negative findings. Second, variability in specimen collection, processing, and storage may have unexpectedly influenced measured protein abundances. Fourth, several proteins known to contribute to ARV disposition were not detected by our proteomic workflow, likely reflecting a combination of factors including the limited quantity of tissue available from rectosigmoid biopsies, the low abundance of some proteins, technical challenges associated with detecting membrane-associated transporters, and the fact that transcript expression does not necessarily correspond to measurable protein abundance. Consequently, absence of detection should not be interpreted as absence of expression or biological importance. An additional limitation is that our proteomic panel did not include several established determinants of ARV disposition, including P-gp, MRP2, OAT1/OAT3, OATP1B1, CES1/2, CatA, and select nucleotide dephosphorylation enzymes. Many of these proteins influence the disposition of ARV classes beyond NRTI; therefore, the present findings should be interpreted as reflecting a subset of intracellular ARV disposition pathways rather than a comprehensive assessment of all mechanisms governing ARV pharmacology. Finally, although proteomics can identify biologically relevant differences in protein abundance, these data alone cannot establish functional consequences for drug disposition or clinical outcomes.

In summary, our findings demonstrate that aging is a major determinant of ARV-disposition related proteomic remodeling within rectosigmoid tissue, influencing transporters, activating enzymes, phosphatases, and, notably, proteins involved in nucleotide homeostasis (**Figure 9**). These effects were further modified by biological sex and occurred within a broader framework of age-associated immune, metabolic, and tissue- specific pathway remodeling. To place these findings in the context of existing literature, **Table 2** compares the observed PBMC and rectosigmoid protein changes with prior reports of aging- and HIV-associated regulation of ARV disposition pathways, highlighting confirmatory, discordant, and potentially novel findings and demonstrating that many of the alterations identified here, particularly within nucleotide homeostasis pathways, have not previously been reported in human tissues. Collectively, our findings identify several candidate mechanisms through which aging may influence ARV disposition at sites relevant to HIV transmission and persistence. As adults over 50 years of age comprise a rapidly growing proportion of both HIV treatment and prevention populations yet remain underrepresented in many pivotal efficacy studies, future clinical pharmacology investigations are needed to determine whether these age-associated molecular changes alter ARV pharmacokinetics, pharmacodynamics, or therapeutic effectiveness. Such studies will be critical for determining whether treatment and prevention strategies optimized in younger adult populations remain equally effective throughout the aging spectrum and whether age-informed therapeutic approaches may ultimately improve HIV outcomes.

**Figure 9.**
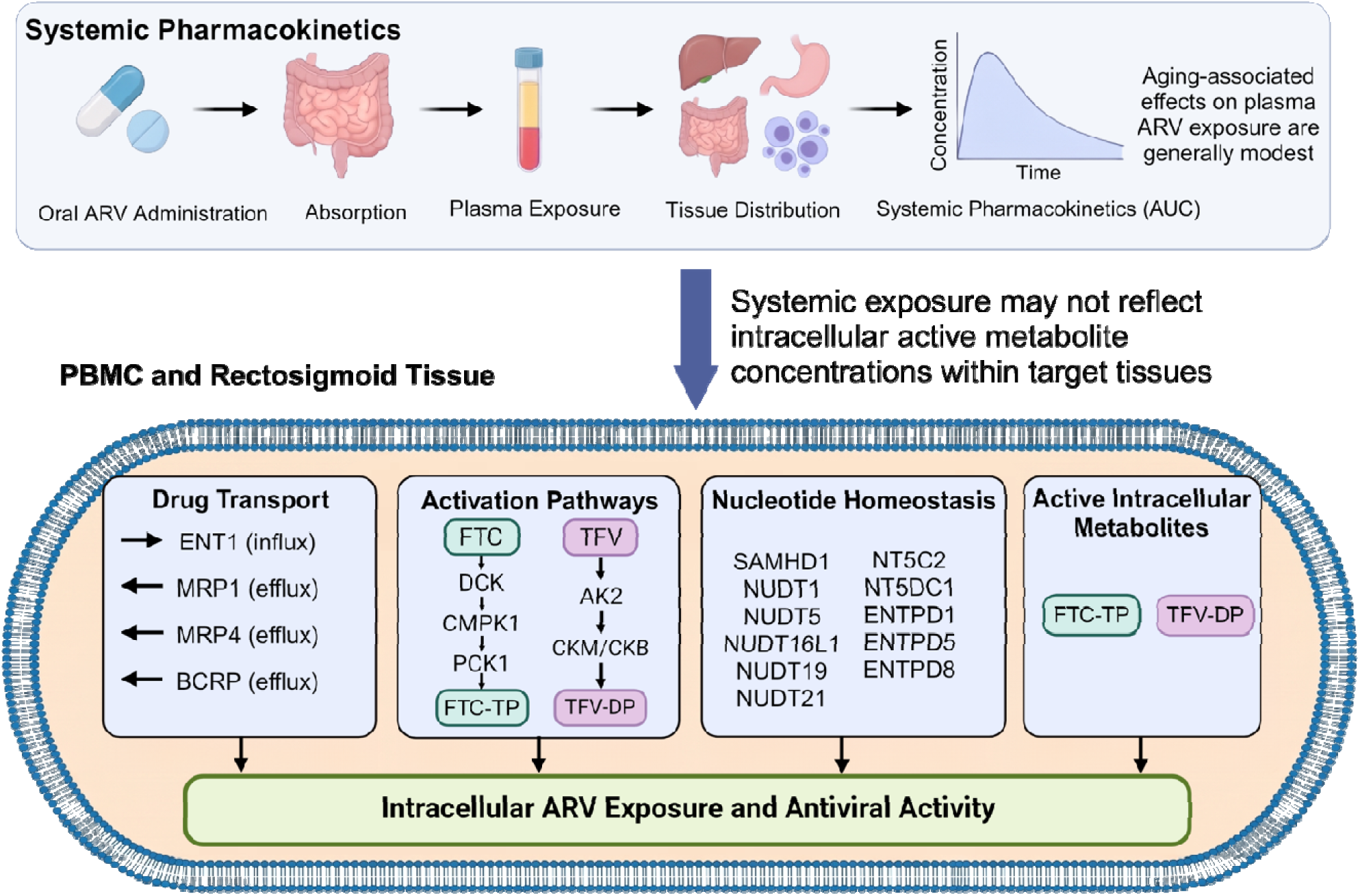
Overview of ARV Disposition and ARV-Related-Proteins of Focus Regulated by Aging and HIV.

**Table 2:** Summary of PBMC and Rectosigmoid Findings in Context of Prior Literature.

| Protein | ARV Relevance | Prior Evidence for Aging/HIV Effects | PBMC Findings | Rectosigmoid Findings | Interpretation | Prior Literature Concordance |
|---|---|---|---|---|---|---|
| **Influx** |  |  |  |  |  |  |
| ENT1 | Nucleoside uptake transporter; contributes to FTC uptake | Among the highest-expressed transporters in colorectal biopsies, with no difference by age, sex or site, though N=12 and mRNA only<sup>50</sup>; no prior PBMC aging data | ↑ 60.1% with aging | No change | Increased nucleoside transport capacity in aging PBMC | Novel (PBMC) |
| **Activation — step 1 (nucleoside/nucleotide kinases)** |  |  |  |  |  |  |
| AK2 | Primary TFV activating kinase (TFV→TFV-MP)<sup>51</sup> | Increased in senescent rat skeletal muscle<sup>35</sup>; no prior human data in any tissue | No change | No change | N/A | Tissue-specific difference from rodent muscle |
| DCK | Initial activation kinase for FTC and related cytidine analogs; also, 3TC and ddC<sup>51</sup> | No age-stratified studies identified in any tissue or species | No change | ↑ 19.8% with aging | Colon-specific regulation of the FTC activation entry step | Novel; discordant between tissues |
| **Activation — step 2 (monophosphate kinases and phosphate donors)** |  |  |  |  |  |  |
| CKM | TFV activation pathway; phosphate donor for TFV-DP formation; also phosphorylates thymidine-analog diphosphates<sup>51</sup> | Activity fell 40–60% in human muscle from donors aged 64–84 vs 24–47<sup>22</sup>; age-dependent nitration<sup>37</sup>; oxidized form unstable with reduced activity<sup>38</sup>; enzyme dysfunction in normal aging brain<sup>23</sup> | No change | ↓ 39.5% with aging | Decreased in rectosigmoid tissue, consistent with established muscle and brain data and the first demonstration outside those tissues | Confirmatory in rectosigmoid tissue |
| CMPK1 | Intermediate FTC/3TC phosphorylation step (monophosphate → diphosphate)<sup>51</sup> | No age- or HIV-stratified data identified in any tissue or species | No change | No change | Unchanged in PBMC and unchanged in rectosigmoid tissue, so the step downstream of the increased colorectal DCK is unclear | Previously unstudied |
| PKM | Step 2 phosphate donor for TFV, with PKLR and CKM<sup>51</sup> | Decreased in senescent rat skeletal muscle<sup>35</sup>; age-dependent nitration of partner glycolytic enzymes<sup>37</sup> | ↓ 34.3% with aging | No change | Decreased in PBMC, consistent with rodent muscle. Unchanged in rectosigmoid tissue, so whether both TFV phosphate donors are aligned is unclear | Confirmatory in PBMC |
| **Activation — step 3 (diphosphate kinases)** |  |  |  |  |  |  |
| PGK1 | Terminal phosphorylation of L-nucleoside analog diphosphates including 3TC/FTC; tenofovir-MP is not a PGK substrate<sup>51</sup> | No age- or HIV-stratified data identified in any tissue or species | No change | No change | Unchanged in both PBMC and rectosigmoid tissue; the first age-stratified measurement of this enzyme in any tissue | Previously unstudied |
| **Catabolism — triphosphate dephosphorylation (ecto-NTPDases)** |  |  |  |  |  |  |
| ENTPD1 | Dephosphorylates nucleotide analog metabolites including TFV-DP; highest activity of NTPDases 1/3/4/5/6/8 toward TFV-DP; with NT5C3 dephosphorylates the active FTC metabolite<sup>52</sup> | Increased in stimulated CD4+ memory T cells of older adults<sup>53</sup> and skin-resident CD4+ T cells, 25.1% vs 12.5%<sup>54</sup>; abundant in colorectal tissue but not previously age-stratified there<sup>52</sup> | No change | ↑ 77.1% with aging | Increased in rectosigmoid tissue, the first age-stratified measurement where rectal PrEP acts | Confirmatory for direction; novel for rectosigmoid tissue; in contrast to previously published T cell data (perhaps because of cell admixture in PBMC) |
| ENTPD5 | ER-luminal UDPase/GDPase; screened against TFV-DP and not the dominant activity<sup>52</sup> | No age- or HIV-stratified data identified | No change | No change | Unchanged in both PBMC and rectosigmoid tissue | Previously unstudied |
| ENTPD8 | Canalicular/apical ecto-ATPase; screened against TFV-DP and not the dominant NTPDase activity<sup>52</sup> | No aging data identified | No change | ↑ 43.6% with aging | Increased in rectosigmoid tissue, but with weaker TFV-DP activity than ENTPD1 | Novel (rectosigmoid) |
| **Nucleotide-homeostasis proteins (NUDT and NT5 families)** |  |  |  |  |  |  |
| NT5C | Dephosphorylates 5′ deoxyribonucleotides; applicable to antiviral nucleoside analogs | No age- or HIV-stratified data identified | No change | No change | Unchanged in both PBMC and rectosigmoid tissue | Previously unstudied |
| NT5C2 | Cytosolic nucleotidase; influences nucleoside analog efficacy through dephosphorylation. Activating mutations confer nucleoside-analog resistance<sup>55</sup> | Known regulator of nucleoside analog response, but no age-stratified studies reported | No change | ↑ 35.7% with aging | Increased in rectosigmoid tissue, but by a margin matching functionally unrelated family members, so not clearly substrate-specific. | Novel (rectosigmoid) |
| NT5DC1 | HAD-superfamily 5′-nucleotidase domain-containing protein; shares the domain only, with no validated nucleotidase activity and no demonstrated NRTI role | No previous HIV- or aging-associated studies | No change | No change | Unchanged in PBMC and unchanged in rectosigmoid tissue; not an NRTI pathway member | Previously unstudied |
| NUDT1 | Oxidized nucleotide sanitizer; main hydrolase of 8-oxo-dGTP. No demonstrated activity against any NRTI triphosphate | Oxidative nucleotide burden (8-oxo-dGTP) increases with aging, but expression versus age is unmapped in normal human tissue | No change | ↑ 32.8% with aging | Increased in rectosigmoid tissue, plausibly reflecting adaptation to age-associated oxidative stress | Novel (rectosigmoid) |
| NUDT5 | ADP-ribose and 8-oxo-dGDP hydrolase; MTH1 (NUDT1) is the major pool sanitizer with little overlapping NUDIX function. No NRTI substrate demonstrated | No previous HIV- or aging-associated studies | No change | ↑ 38.4% with aging | Unchanged in PBMC, increased in rectosigmoid tissue, matching the coordinated rise across this category in mucosa | Novel (rectosigmoid); discordant between tissues |
| NUDT16L1 | TIRR, a 53BP1-binding DNA repair regulator. Catalytically inactive relative to NUDT16; no demonstrated nucleotide-pool or NRTI role | No previous HIV- or aging-associated studies | ↓ 25.3% in PLWH | ↓ 26.6% with aging | A decrease of comparable magnitude in each tissue. Reproducible, but suggests an effect on the DNA damage response rather than on nucleotide or NRTI metabolism | Novel; the only finding of concordance between tissues |
| NUDT19 | Peroxisomal CoA diphosphatase (acyl-CoA/CoA turnover); not a nucleoside-analog enzyme | No previous HIV- or aging-associated studies | ↓ 50.0% in PLWH | No change | Halved in PBMC, the largest single decrease across both tissues, but in peroxisomal CoA handling rather than an NRTI pathway. | Novel (PBMC) |
| NUDT21 | CFIm25, cleavage factor Im subunit; RNA 3′-end processing and alternative polyadenylation. Not a nucleotide hydrolase | Studied in senescence biology via 3′UTR shortening, but no ARV-relevant aging data available | No change | ↑ 36.0% with aging | Unchanged in PBMC, increased in rectosigmoid tissue in line with its other proteins in this category; supports broader transcriptomic remodeling rather than an ARV-specific effect | Novel; discordant between tissues |
| **Competing dNTP pool** |  |  |  |  |  |  |
| SAMHD1 | Regulates dNTP pools affecting NRTI activity; hydrolyzes NRTI-TPs far more slowly than dNTPs | Thymidine-analog-selective effect on NRTI potency<sup>56</sup>; markedly increased in quiescent cells with falling dNTP pools<sup>57</sup> | No change | No change | Unchanged in PBMC and unchanged in rectosigmoid tissue | Expected increase not observed |
| **Efflux** |  |  |  |  |  |  |
| BCRP | Efflux transporter of broad ARV classes<sup>51</sup> | Hepatic BCRP protein decreases with aging while mRNA is unrelated to age<sup>20</sup>; duodenal BCRP increases with aging<sup>21</sup>; expressed in colorectal biopsies but not age-stratified there<sup>50</sup> | No change | ↑ 115.6% with aging | Increased in rectosigmoid tissue and was the largest effect in this study | Confirmatory for duodenum; opposite to liver confirming that BCRP aging-related effects are region-dependent |
| MRP1 | Efflux transporter of broad ARV classes<sup>51</sup> | Age-dependent increase in human duodenum<sup>21</sup>; measured in PBMC but not age-stratified<sup>58</sup>; surveyed in colorectal tissue without an age effect<sup>50</sup> | No change | No change | Unchanged in PBMC and unchanged in rectosigmoid tissue | In contrast to MRP4 findings suggests substrate-specific rather |
| MRP4 | Major TFV/TFV-DP efflux transporter; principal export route for NRTI monophosphates<sup>51</sup> | Age-dependent increase in human duodenum<sup>21</sup>; immature relative to OAT1/OAT3 in developing rat kidney<sup>59</sup>; measured in PBMC but never age-stratified<sup>58</sup>; surveyed in colorectal tissue without an age effect<sup>50</sup> | ↑ 107.8% in PLWH | No change | Increased in PBMC (second largest effect in this study) | Novel (PBMC); consistent with duodenum |

↑ / ↓ denote the direction of change.

**a)** This work was supported by the National Institutes of Health, National Institute of Allergy and Infectious Diseases (R56AI161030), National Institute on Drug Abuse (R01DA052859, U01DA058527, R01DA063076, and R01DA066140), National Institute of Neurological Disorders and Stroke (K00NS118713), National Institute for General Medical Sciences (T32GM066691, T32GM008763, T32GM144272), the Johns Hopkins University Center for AIDS Research (P30AI094189), and the Center for AIDS Research at Emory University (P30AI050409). The content is solely the responsibility of the authors and does not necessarily represent the official views of the National Institutes of Health.
**b)** This work was conducted in partial fulfillment of the requirements for the Ph.D. degree from The Johns Hopkins University in 2024. The work described in this manuscript was performed while authors, Colten Eberhard and Hannah Wilkins, pursued doctoral studies at The Johns Hopkins University School of Medicine, prior to the authors’ employment or affiliation with their respective current companies.
**c)** Please contact Dionna W. Williams, 1510 Clifton Road, Atlanta, GA 30322, for reprint requests

## Nonstandard abbreviations

AK2: Adenylate kinase 2
ARV: Antiretroviral
ART: Antiretroviral therapy
BCRP: Breast cancer resistance protein (ABCG2)
BIC: Bictegravir
CKB: Creatine kinase, brain-type
CKM: Creatine kinase, muscle-type
CMPK1: Cytidine monophosphate kinase 1
DCK: Deoxycytidine kinase
ENT1: Equilibrative nucleoside transporter 1 (SLC29A1)
ENTPD1: Ectonucleoside triphosphate diphosphohydrolase 1
ENTPD5: Ectonucleoside triphosphate diphosphohydrolase 5
ENTPD8: Ectonucleoside triphosphate diphosphohydrolase 8
FTC: Emtricitabine
GI: Gastrointestinal
LC-MS: Liquid chromatography-mass spectrometry
MMTS: Methyl methanethiosulfonate
MRP1: Multidrug resistance-associated protein 1 (ABCC1)
MRP4: Multidrug resistance-associated protein 4 (ABCC4)
NRTI: Nucleos(t)ide reverse transcriptase inhibitor
NT5C2: Cytosolic 5′-nucleotidase II
NT5DC1: 5′-nucleotidase domain-containing protein 1
NUDT1: Nudix hydrolase 1
NUDT5: Nudix hydrolase 5
NUDT16L1: Nudix hydrolase 16-like 1
NUDT19: Nudix hydrolase 19
NUDT21: Nudix hydrolase 21
PBMC: Peripheral blood mononuclear cell
PGK1: Phosphoglycerate kinase 1
PKM: Pyruvate kinase M
PLWH: People living with HIV
PrEP: HIV pre-exposure prophylaxis
SAMHD1: SAM and HD domain-containing protein 1
SDS: Sodium dodecyl sulfate
TAF: Tenofovir alafenamide
TDF: Tenofovir disoproxil fumarate
TFV: Tenofovir
TFV-DP: Tenofovir diphosphate

## Acknowledgments

We also thank Kate Marych, RN for her outstanding leadership and dedication as senior research nurse for this study. Her contributions to participant recruitment, clinical coordination, specimen collection, and participant care were essential to the successful completion of this work. Additionally, we are grateful for scholarly discussion and intellectual contribution with current and past members of the Williams laboratory. We also would like to thank the Clinical Pharmacology Analytical Laboratory at The Johns Hopkins University School of Medicine for performing TFV and TFV-DP quantitation using our processed samples.

## Authorship Contributions

Participated in research design: H.N.W., R.Z., N.N.B., C.W.H., B.C.O.

Conducted the clinical study: E.J.F, J.H., R.Z., E.G., C.W.H.

Conducted experiments: H.N.W.,C.D.E., B.C.O., R.Z.

Contributed new reagents or analytic tools: H.N.W., B.C.O., C.W.H.

Performed data analysis: H.N.W., C.J.W., R.Z.

Wrote or contributed to the writing of the manuscript: C.J.W., H.N.W., D.W.W, C.W.H., B.C.O.

---

**Figure S1.**
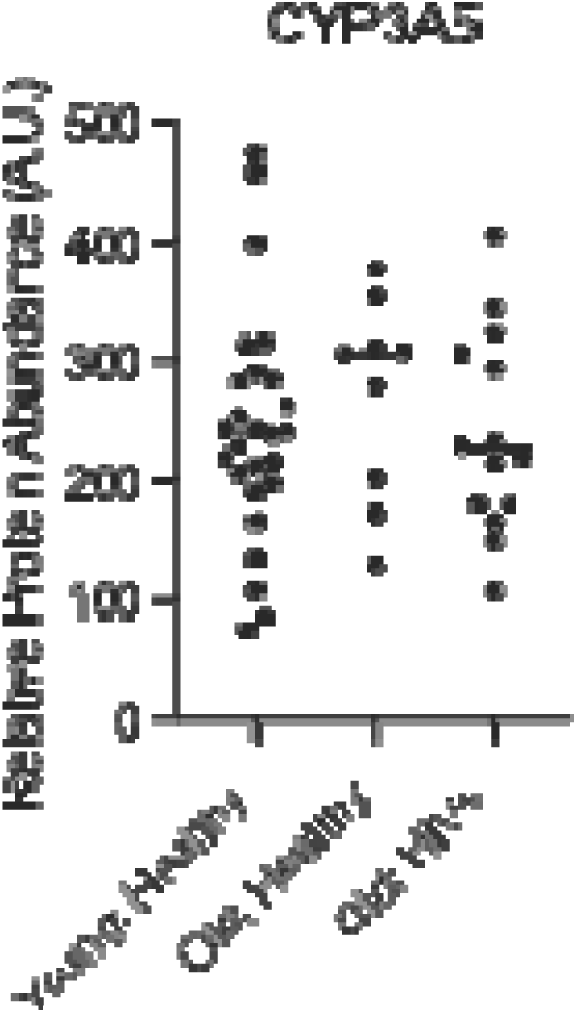
CYP3A5 is unchanged in by aging and HIV in human rectosigmoid tissue. Human rectosigmoid tissues were isolated from young and old, HIV negative participants as well as old, HIV positive participants for measurement of CYP3A5 concentrations. All participants in the old, HIV positive cohort were on a TFV-based ART regimen for treatment of HIV. Concentrations were measured by Proteomic Ruler. A minimum of three independent experiments were performed for rectosigmoid tissue LC-MS proteomics with two LC-MS injection replicates except for the old, HIV negative group which was limited to two independent experiments (represented as individual dots). D’Agostino & Pearson, Anderson-Darling, Shapiro-Wilk, and Kolmogorov- Smirnov test were performed to assess. normality and lognormality. Data represented as mean ± standard deviation. Statistical analysis was performed for normally distributed data using Brown–Forsythe and Welch ANOVA by GraphPad software. *p < 0.05, **p < 0.01, ***p < 0.001, and **** p < 0.0001.

## Data Deposition

All vendor raw intermediate processed data was deposited at the ProteomeXchange partner repository MASSIVE (massive.ucsd.edu) as accession PXD061862 and is permanently publicly available for reader and reviewer access.

